# Counterfactual reasoning underlies the learning of priors in decision making

**DOI:** 10.1101/227421

**Authors:** Ariel Zylberberg, Daniel M Wolpert, Michael N Shadlen

## Abstract

Accurate decisions require knowledge of prior probabilities (e.g., prevalence or base rate) but it is unclear how prior probability is learned in the absence of a teacher. We hypothesized that humans could learn base rates from experience making decisions, even without feedback. Participants made difficult decisions about the direction of dynamic random dot motion. For each block of 15-42 trials, the base rate favored left or right by a different amount. Participants were not informed of the base rate, yet they gradually biased their choices and thereby increased accuracy and confidence in their decisions. They achieved this by updating knowledge of base rate after each decision, using a counterfactual representation of confidence that simulates a neutral prior. The strategy is consistent with Bayesian updating of belief and suggests that humans represent both true confidence, that incorporates the evolving belief of the prior, and counterfactual confidence that discounts the prior.

## Introduction

Accurate decision making relies on both evidence bearing on the choice at hand and prior knowledge about the statistical regularities bearing on the possible options. In some instances we learn these regularities through education (e.g., disease prevalence based on decisions of experts), but more often such knowledge is acquired over time through decisions we make ourselves. This poses a problem because absent an omniscient teacher, our decisions can be inaccurate, which limits our ability to update our beliefs. The problem is compounded because the decisions we make may be affected by our evolving belief of the statistical regularities, thereby biasing our decisions, which could in turn affect how we update our belief about the regularities.

As an example, consider you are working on a production line determining whether tomatoes arriving one at a time on a conveyor belt are ripe, and ready to ship, or unripe and need to be held back. Some are obviously red and ripe and others are clearly green and unripe but there are others on the border which will be harder to judge. Each crate comes from a different supplier and will likely have a different proportion of unripe tomatoes. If you knew each crate’s proportion of unripe tomatoes (the base rate) it would help you to sort the tomatoes. This is an example in which one is learning a prior (base rate) without a teacher or confirmation about ground truth.

One possibility is that one would ignore base rate in this setting, but that would be unwise when confronted with an ambiguous tomato. It makes more sense to estimate the base rate from one’s experience. One could simply use the proportion of decisions of unripe as the estimate of base rate. However, this would give equal weighting to tomatoes judged as clearly unripe and those on the borderline. An alternative might be to use a measure of confidence in determining the base rate, but this invites another challenge. If one uses the base rate to decide ripeness, this will affect the tally and potentially bias the estimate of the base rate itself.

This simple example illustrates the complexity of doing inference in a world in which one is simultaneously learning a model and applying it. It arises in medical decision making (Medin & Edelson, 1988; Goldberg, 1970), weather prediction (Knowlton et al., 1994; Yang & Shadlen, 2007), and other inference problems that benefit from experience but for which feedback about ground truth is unavailable on a useful time scale. It has been shown that people are able to estimate base rates from a sequence of observations (Phillips & Edwards, 1966; Estes, 1972; Peterson & Beach, 1967) to develop a bias that serves as prior knowledge in subsequent interactions with the environment (Anderson & Carpenter, 2006; Manis et al., 1980), and even use these observations to infer changes in the state of the environment (Summerfield et al., 2011; Behrens et al., 2007; Meyniel et al., 2015; Purcell & Kiani, 2016; Nassar et al., 2010; Rapoport, 1964; Carpenter & Williams, 1995). In these cases, the observations that inform the prior are clearly discernible (e.g., target reached / not-reached) or are accompanied by explicit feedback. For example, if the sorter were to taste each tomato the ground truth could be known, but there would be no tomatoes shipped. We hypothesized that in the absence of explicit feedback, decision confidence guides the acquisition of prior probability and does so in accordance with Bayesian updating. We build on recent progress in understanding of confidence in simple perceptual decisions.

To study the role of confidence in the acquisition of prior probability, we designed a task in which human participants made a sequence of binary decisions in the presence of a concealed base rate that favored one of the alternatives. The decisions involved judging the direction of motion of a set of randomly moving dots, which were made without feedback. The base rate was constant within a block of trials but randomly varied from one block to another. Crucially, the base rate was not known to the participant. As the participants made more decisions, the influence of base rate on choice and confidence increased, which was reflected both in the decision about the direction of motion and in an explicit report about the bias of the block. A bounded evidence accumulation model explained the decisions about motion by incorporating an estimate of the base rate in the accumulation. In turn, a probability distribution over base rates was updated based on the likelihood that the motion was rightward or leftward—what we term counterfactual posterior probability—under the fictitious supposition that the alternatives are equally likely. The model predicted the dynamics of belief about the direction bias over the block. The findings expose a role for counterfactual confidence in belief updating, suggesting that the brain maintains probabilistic representations over decision hierarchies and time scales: direction over one trial and bias over many trials. Further, these probabilities are accessible for explicit reporting.

## Results

Three participants were presented with a dynamic display of random dots of variable duration and had to decide whether the net direction of motion was rightward or leftward. Within a block of 15 to 42 trials, one direction of motion (left or right) was more likely, but which direction was more likely (and by how much) was unknown to the participant. The difficulty of the decision was controlled by three factors: strength of motion, stimulus duration, and bias strength (i.e., base rate). Motion strength was controlled by the probability (termed motion coherence, *c*) that a dot is displaced in motion as opposed to randomly. Stimulus duration was sampled from a truncated exponential distribution (range: [0.1,0.9] s). Bias strength was controlled by the probability that the motion direction would be rightward, termed *B*, which was selected randomly on each block from six possible values ranging from 0% to 100% in steps of 20%. Participants knew the possible values of *B*, and that they were equally likely, but were not told which one applied to the current block.

Participants made three responses in each trial. They first reported the perceived direction of motion and the confidence that this decision was correct (Figure 1A, choice and confidence report). They then reported whether they considered the block to have a right or left bias, and the confidence in this judgment (Figure 1A, Belief report). To avoid confusion, we refer to this type of confidence as ‘belief’: an estimate of the probability that the block has a rightward bias (scale 0 to 1). Participants received no feedback about the accuracy of their decisions during a block of trials. Only after completing a block, were they told which direction was the most probable, the strength of the bias in this direction (either 60, 80 or 100%) and the proportion of trials in which they responded correctly (Figure 1B).

**Figure 1.**
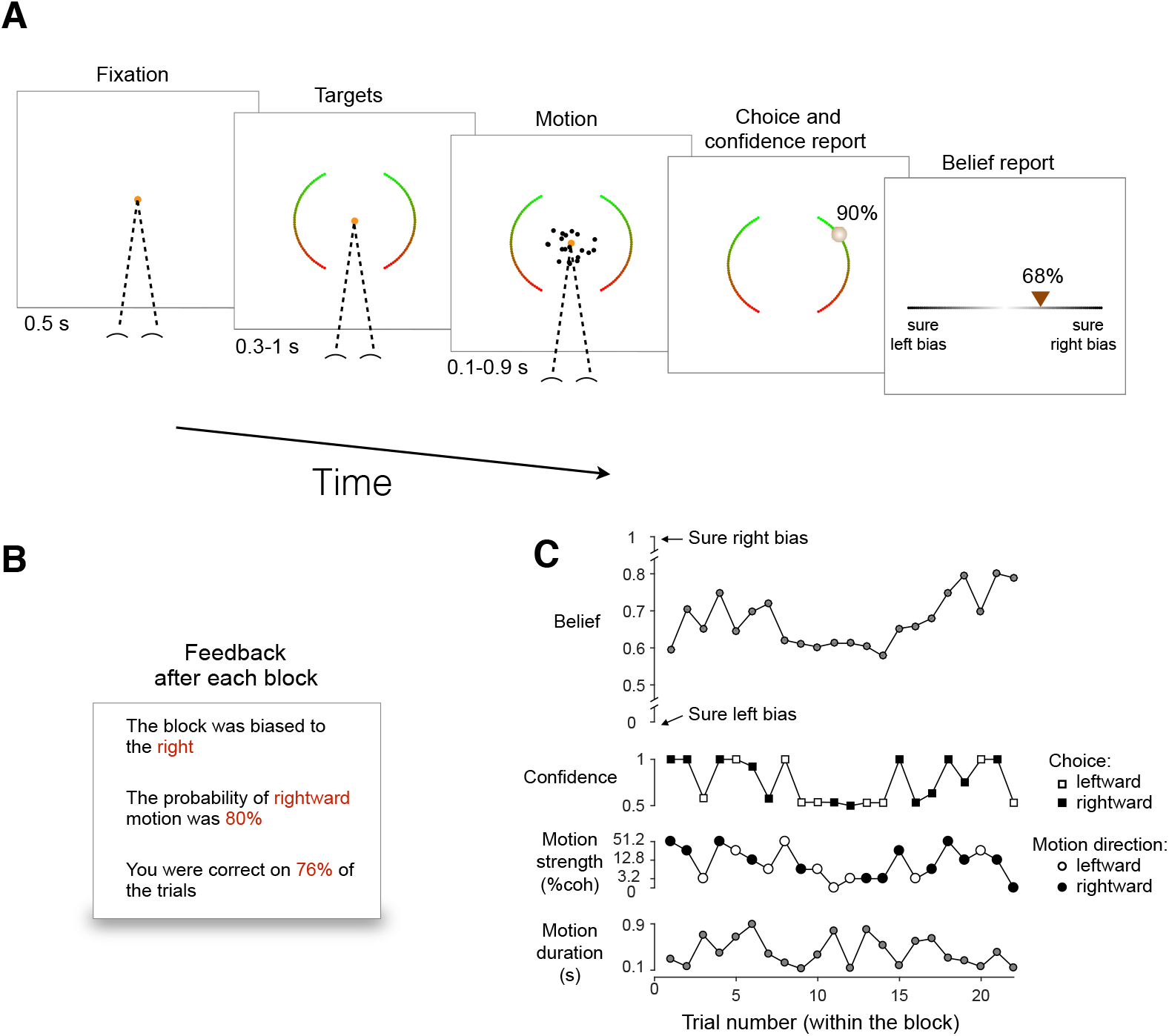
Motion discrimination task. Each block of 15 to 42 trials was assigned one of 6 possible base rates: the prior probability the motion is rightward. In all blocks the subject discriminated the direction of random dot motion. No feedback about individual decisions or the base rate was provided until the end of the block. (**A**) Sequence of events within a trial. After fixation and a random delay, random dot motion was displayed for 0.1-0.9 s (truncated exponential). Subjects then positioned a cursor on the left or right arcs to indicate both choice (left vs. right) and confidence (100% certainty top to guessing bottom). After the motion decision, the subjects reported whether they believed that the block had a rightward or leftward bias (placing the cursor in the left or right half of the line), together with the confidence in this belief (from unsure at the center to certainty at the extremes). (**B**) Example of the feedback display provided at the end of a block. (**C**) Example of a sequence of trials within a block. Lower two graphs show variables controlled by the computer: motion strength, direction and duration. Upper two graphs show the subject’s reports: direction choice, confidence that the choice was correct, and belief that the base rate favored rightward. A movie of the task is shown in Movie S1.

Figure 1C shows a typical sequence of events and reports that transpire in a single block in the experiment. The bottom two rows show the particular sequence of 22 trials, each associated with motion stimulus of some strength, direction and duration. The upper two rows show the corresponding behavioral reports: direction choice, confidence in the choice, and belief about bias of the block. In this block, the participant was correct on most of the trials, with a confidence that was strongly correlated with motion strength. At the beginning of the block, the belief was in the region of high uncertainty (~0.5) and increased later in the block. As a glimpse into what we will appreciate in greater detail later, it can be seen that decisions made with high confidence are usually followed by larger changes in belief than decisions made with low confidence. This is evident for trials 9 to 14. The example provides an intuition for the inference problem the participant confronts on each short block of 15 to 42 trials.

We next describe the main effect of the base rate on the direction choices, the confidence in these choices and the belief that the block is biased to the left or right. We then develop a theory to explain the way the direction decisions inform belief and the way belief biases those choices. Finally, we use this theory to predict the time course (evolution) of this belief. We contrast this theory with alternatives.

### Effect of base rate on choice, confidence and belief

Throughout each block, choices were governed by the strength and direction of random dot motion. Figure 2A shows the proportion of rightward choices as a function of stimulus strength, combining data from all trials sharing the same base rate (color). When the base rate strongly favored rightward or leftward (1 or 0, respectively) nearly all of the choices were consistent with the base rate. At the intermediate base rates, the shift was less pronounced. To capture the effect in a model free way, we performed logistic regression (solid curves) and estimated the choice bias (Eq. 17, Methods). As shown by the inset, the subjects clearly internalized the base rate during the block (Eq. 17; *p* < 10^−8^; likelihood-ratio test; *H*_0_: all *β*_3_ = 0). That is, for the same motion strength, subjects were more likely to choose the direction consistent with the base rate of the block. We show combined data from all subjects (individual subjects are presented in Figure S1). It is clear from these observations that the subjects acquire knowledge about the base rate of the block, despite the absence of feedback about whether their decisions were correct. Knowledge of the base rate should improve the performance on the direction task. This is clearly supported by Figure 2B, which shows accuracy as a function of motion strength (|*c*|), and groups base rates of similar strength. We adduce from these observations that participants incorporated knowledge of the base rate to bias and improve their decisions.

**Figure 2.**
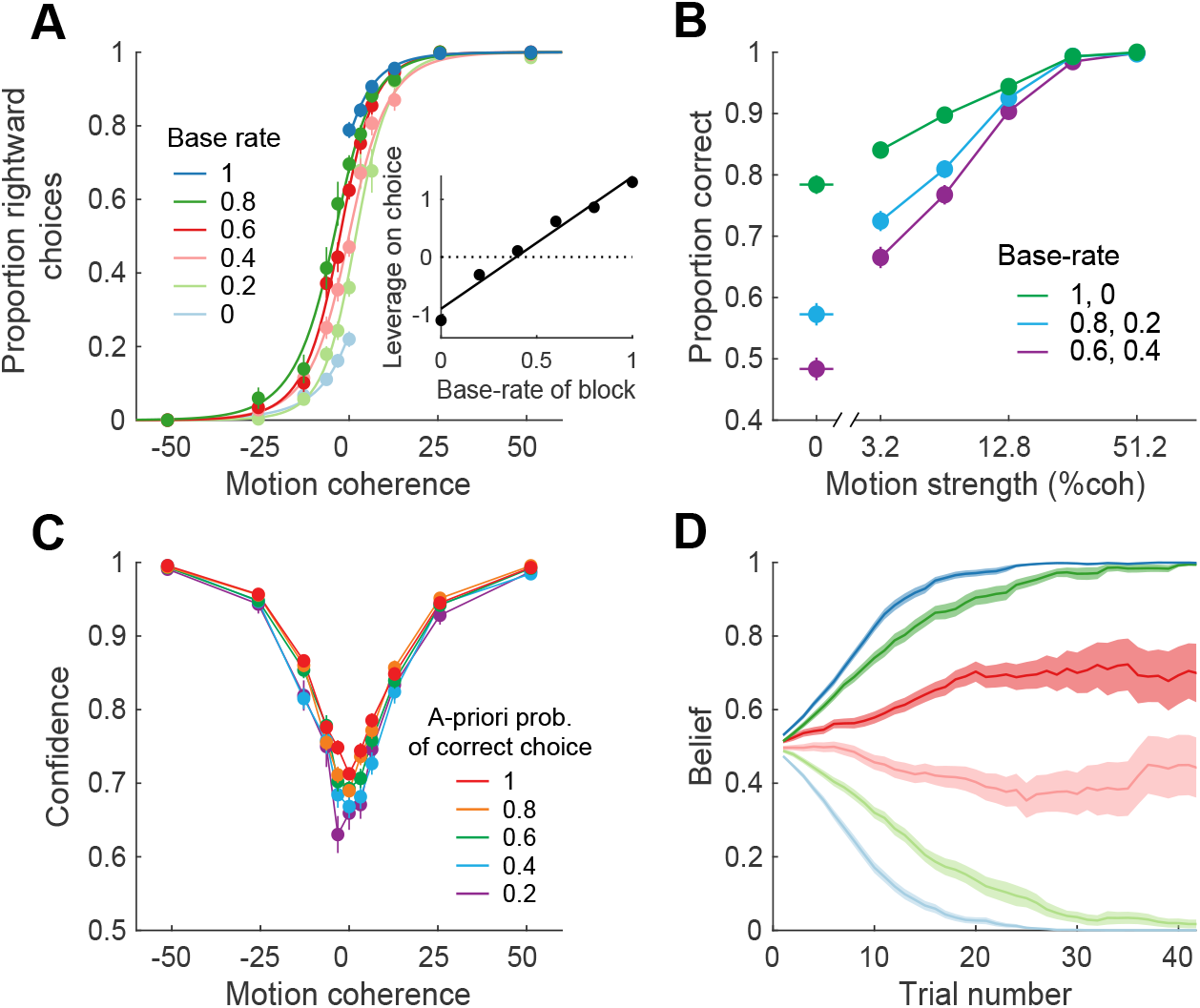
Behavior was influenced by the base rate. (**A**) Proportion of trials on which the participants reported rightward motion, as a function of motion coherence. Data (symbols) are shown separately for the six levels of base rate, from 0 (leftward was the correct choice for all trials in the block) to 1 (all rightward), combining all stimulus durations and trials in the block. The solid lines are fits of a logistic regression model (Eq. 17). The shift in the psychometric functions indicate that participants choices were influenced by the base rate. Inset shows the magnitude of the bias (*β*_3_, equation 17) against the actual base rate (error bars are s.e.; most are smaller than the points; solid line is least-squares fit). (**B**) Effect of motion strength and base rate on choice accuracy (same data as in A).The base rates (color) are combined by degree of informativeness. (**C**) Average confidence reported on correct trials as a function of the motion coherence. The *a priori* probability of correct (color) is an expression of the base rate relative to the choice that is made (see text). Error bars represent s.e. across trials. (**D**) Average belief as a function of trial number within a block for the six different biases. Same color convention as in Panel A. A belief of 0 or 1 indicates full certainty that the block was biased to the left or right, respectively. Shading indicate s.e.m. Data are combined across all participants. The same analyses for each participant are shown in Figure S1, and the distributions of confidence and belief for each subject are shown in Figure S2.

Subjects furnished two additional reports, confidence and belief, which indicate that they formed an impression of the prior probability about direction over the course of a block. The confidence reports associated with each decision were clearly influenced by the base rate. Fig 2C shows the confidence ratings for correct choices split by the *a priori* probability that the direction of motion supports the choice that the participant made (correct choices only). For example, the *a priori* probability of 0.8 groups together right choices with rightward motion (positive coherence) in blocks with base rate of 0.8 and left choices with leftward motion (negative coherence) in blocks with base rate of 0.2. Two features of the confidence ratings stand out. The subjects were least confident when the motion was weak (coherence near zero; Eq. 18; *p* < 10^−8^, t-test, *H*_0_ : *β*_1_ = 0). Moreover, confidence was higher when the base rate was more informative (Eq. 18; *p* < 10^−8^, t-test, *H*_0_ : *β*_2_ = 0). This is important, because it implies that subjects were not merely choosing one direction more often (e.g., out of habit) but that they incorporated knowledge of the base rate to reduce uncertainty in the decision.

The second report was the belief that the base rate of the block favored right or left. This “belief” evolved gradually during the course of the block (Fig. 2D). Note that the belief is not an estimate of the base rate, as one can be fully confident in a weak bias, but its evolution was more rapid on average when the base rate was more informative (Fig. 2D, blue curves). We will attempt to explain the evolution of this bias by developing a theory of the two way interaction between bias and choice—that is, the effect of bias on each decision and the effect of each decision on the estimate of the base rate of the block.

### Hierarchical Bayesian model

We developed a hierarchical Bayesian model in which subjects maintain a probability distribution over the base rate, *p*(*B*), within a block and use this knowledge to influence both their choice and confidence within a trial. As we will see, the optimal way to update *p*(*B*) is to use a counterfactual form of confidence—the probability that a choice rendered on the evidence would be correct if the base rate were unbiased (i.e., *B* = 0.5). We develop this idea in Figure 3 and provide a mathematical derivation in Methods.

**Figure 3.**
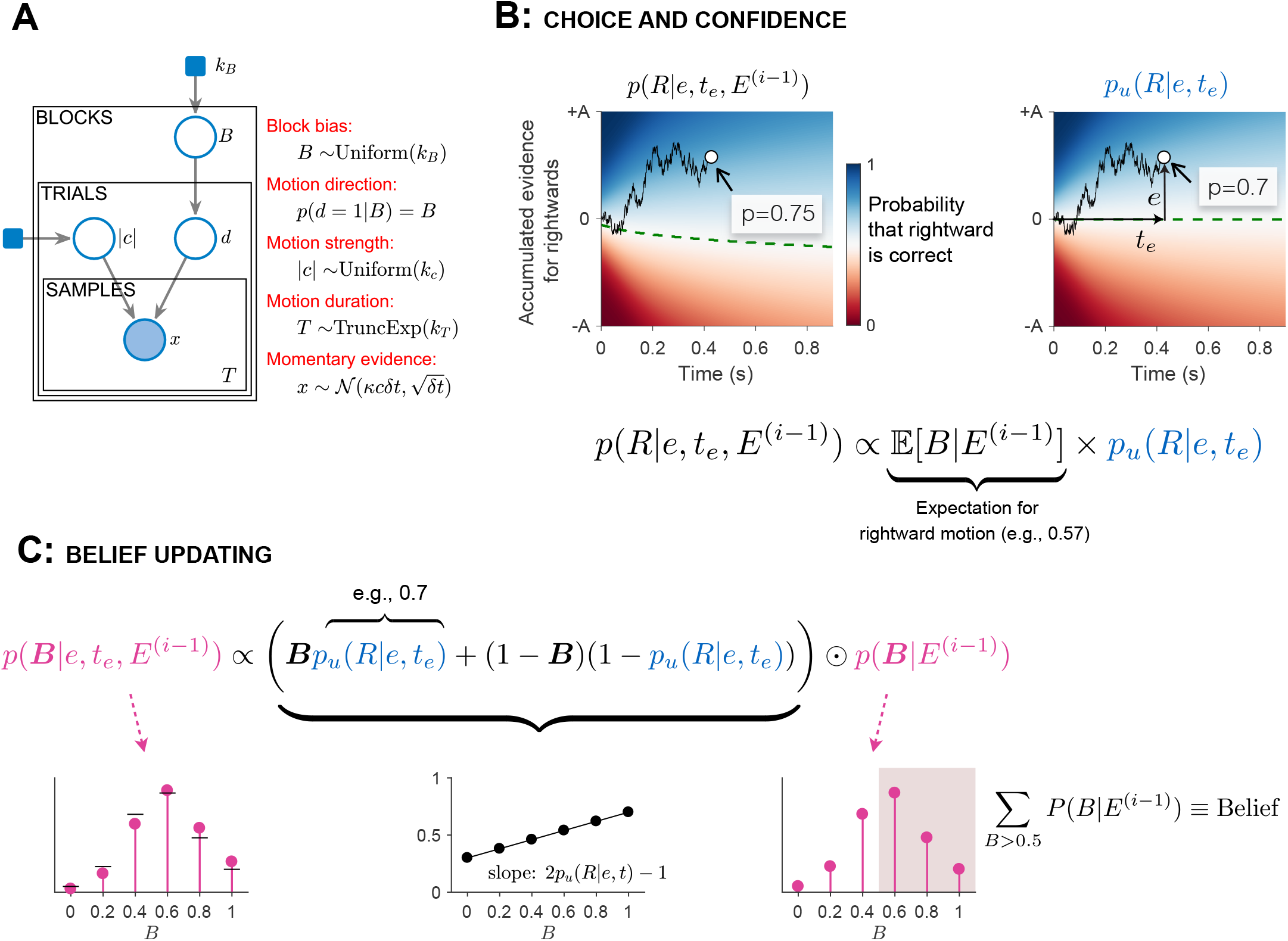
Bayesian model. (**A**) Graphical representation of the hierarchy of causes that give rise to a sample of momentary motion evidence. The shaded areas are observed variables; the unshaded areas are unobserved variables; and the filled squares are fixed hyperparameters. The block bias is sampled from a uniform distribution over six possible values (parameterized by *k_B_*). B is the block’s base rate which sets the prior probability over motion direction (*d*) for the entire block. The motion strength (|*c*|) is sampled on each trial from a uniform distribution over six possible values (parameterized by *k_c_*), independent of the block’s base rate. In the experiment, |*c*| and *d* set the probability that a dot plotted at time *t* will be replotted at time *t*+40ms toward the right or left target. The duration of motion (*T*) was sampled from a truncated exponential distribution. In the model, it is assumed that the momentary evidence follows a Gaussian distribution with a fixed variance (Δ*t*) and a mean that depends on |*c*| and d. (**B**) Bounded evidence-accumulation model of choice and confidence. The decision is made when the accumulation of momentary evidence (*e*) reaches a bound (±*A*) or the stimulus is curtailed, as in the sample trace. The two color maps show the probability that rightward would be the correct choice based on accumulated evidence *e* at time *t_e_*, under different assumptions about the base rate: (*left*) expectation of the base rate is 0.57 in favor of rightward; (*right*) based rate is unbiased (0.5). The dashed green line marks the decision boundary (*p*(*R*) = 0.5). We refer to the right map (*p_u_* (*R*|*e, t_e_*)), as the counterfactual posterior probability of rightward. The left map is formed by multiplying the counterfactual posterior by the expectation of the base rate, given the evidence from the previous trials (*E*^(*i*−1)^). The normalization constant assures that the posterior over direction sums to unity. (**C**) Belief updating. To update *p*(*B*|·), the distribution over base rate from the previous trial (shown on the right) is pointwise multiplied (⨀) by the expression inside the parenthesis. This expression is a linear function of *B*, with a slope given by the counterfactual posterior (center plot). The left panel shows the updated distribution after the pointwise multiplication and normalization. For reference, the horizontal black lines indicate the distribution over base rates from the previous trial. The normalization constant ensures that *p*(*B*) sums to unity. The belief that the block is biased in favor of rightward is given by the sum of *p*(*B*) for *B* > 0.5 (shading).

Figure 3A shows the hierarchy of causes that give rise to a sample of motion evidence, *x*. All trials are affected by the base rate, *B*, assigned to the block, and *B* itself is sampled from a uniform distribution. The bias in the block establishes the prior probability of direction (*d* = *sign*(*c*), i.e. R or L) of motion on each trial. The strength of motion (|*c*|) and the stimulus duration (*T*) are determined probabilistically for all trials (independent of block). The direction and strength of motion specify the stationary stochastic process comprising samples of evidence, *x*. We assume that *x* is a sample from a Gaussian distribution with mean=*κc*Δ*t* and variance equal to the sample period (Δ*t*). The parameter *κ* reflects the signal-to-noise in the accumulation process. To make a decision, the brain accumulates samples until either the accumulated evidence (*e*) reaches a threshold at ±*A* or until the end of the motion stimulus (Fig. 3B). The time of this last sample is denoted *t_e_*. If leftward and rightward are equally likely, the decision about direction should be determined by the sign of *e*, and the probability that the motion is rightward is determined by *e* and *t_e_* (Kiani & Shadlen, 2009). The heat map (Fig. 3B right) shows the probability that the direction is rightward for all possible combinations of *e* and *t_e_*: *p_u_* (*R*|*e*, *t_e_*). The example trial (open circle) would have led to a rightward choice with probability 0.7 of being correct. The top half of the map thus provides a lookup table for confidence in a rightward choice. Leftward choices would arise when the evidence (*e*) is less than 0, and the probability that this choice is correct is 1 minus the values displayed, that is, the top half of the map reflected vertically across the horizontal green line. These statements about direction choice, probability of left/right and confidence apply only if leftward and rightward motion are equally probable—hence the subscript, *u* (for unbiased), in *p_u_*(*R*|*e, t_e_*). This condition does not occur in our experiment, but the mapping plays a role. We refer to the mapping on the right as a counterfactual posterior or counterfactual confidence.

In the experiment, one direction within a block is always more likely than the other, and this affects the probability that the direction is right (or left) given {*e, t_e_*}. For example if the base rate, *B*, favors rightward, it might give rise to the map in Figure 3B (left). Notice that the decision criterion (dashed green line) dips to negative values of *e*. The mapping to confidence is altered as well. The same amount of evidence leading to a right choice (circle) now corresponds to 0.75 probability of being correct. The calculation supporting this map factors neatly into the expectation of the base rate multiplied by the counterfactual posterior—the probability that the direction would be rightward if the the two directions were equally likely. The expectation of the base rate is informed by knowledge obtained on the previous trials in the block, represented by *E*^(*i*−1)^ ≡ {*e*_1..*i*−1_, *t*_*e*,1..*i*−1_}. In the example, the expectation of B is 0.57. The confidence is 0.57 × 0.7 divided by the sum of this term plus (1 − 0.57) × (1 − 0.7), where the last product is *p*(*d* = *L*|*E*^(*i*−1)^)*p_u_*(*L*|*e, t_e_*). The arithmetic yields approximately 0.75, which is also the posterior probability of right. The intuition is straightforward: given knowledge of the base rate through the previous trial, use its expectation to adjust the decision criterion and confidence based on the evidence and termination time on the present trial.

The remaining question is how knowledge of the base rate is updated. According to the hierarchical Bayesian model, the subject begins the block with a prior over the 6 possible base rates, *p*_0_(*B*). The true prior is uniform, but we allow for the possibility that subjects do not internalize this correctly. Panel C shows how these values are updated. As the graphical model (panel A) makes clear, inference about *B* is arbitrated solely by the direction of motion, *d*. Key to the update is that the subject should not use the estimate of direction that they report but the probability of each direction under a neutral prior (*B* = 0.5), in other words the counterfactual posterior. This yields the update rule illustrated in Figure 3C, which depends on two components which change across trials: the posterior for rightward (and its complement for leftward) and the current estimate of the distribution of *B*. The use of the posterior can be visualized as a line (center, panel C) that is then point-wise multiplied by the current *p*(*B*) (right, panel C). The six probabilities are scaled to sum to unity (left, panel C). Note that the updated distribution after one trial becomes the initial distribution for the next one (i.e., *p*(*B*|*E*^(*i*)^) ≡ *p*(*B*|*e, t_e_, E*^(*i*−1)^)). Again, the full derivation of the expressions is in Methods.

To appreciate why the Bayesian solution uses the counterfactual posterior to update *p*(*B*), consider the following example. Suppose that the participant has acquired a slight bias for rightward, and is then presented with a sequence of stimuli of 0% coherence. The participant will tend to report rightward more often, because the bias exerts a stronger influence when the evidence is weak. Therefore, if true posterior (confidence) is used to update the bias, the bias would tend to increase until the decision maker is certain that the block contains a rightward bias, even though the sensory evidence is ambiguous. As formalized by the expression in Figure 3C (and in Methods, equations 6 to 8) the correct approach is to update the belief based on the likelihood that the evidence (*e, t_e_*) had been obtained in a trial with right or left motion, or, equivalently, the confidence that the subject would have under a neutral (i.e., uninformative) prior. In short, the counterfactual posterior circumvents the problem of a self-reinforcing prior, what might be thought of as double counting.

This completes both parts of the theory: (*i*) how to incorporate one’s estimate of bias into the choice and confidence about direction, and (*ii*) how to update the estimate of bias (including the confidence in this estimate) based on this experience.

### Fits of the Bayesian model to choice and confidence

Our main hypothesis is that participants modify their bias according to the counterfactual confidence they have in their decision. To test this hypothesis, we use the Bayesian model described in the previous section to fit each subject’s choices and confidence reports. We compared this model to two alternative models, one in which participants update their bias based only on the frequency of left and right choices (i.e., ignoring confidence), and another in which participants used confidence (instead of its counterfactual) to update the bias. After comparing the goodness-of-fit of the three models for the motion direction reports, we use the best-fitting model to predict—without additional degrees of freedom—the explicit reports of the belief which were not used in the model fitting.

We used the sequence of stimuli (motion coherence and duration) on each trial to fit the choice and confidence reports. The model constructs a hidden “latent” representation of the subject’s knowledge of the base rate, *p*(*B*), as it it evolves with each trial. As shown in Figure 4A subjects’ choice accuracy was explained by the Bayesian model. The points, which are identical to those in Figure 2B, combine data over the entire block, both directions, and all stimulus durations. The more informative the base rate, the more it can be relied upon to improve accuracy. The model explains quantitatively the degree to which the base rate was learned and incorporated by the participants (see Fig. S3 for individual subjects). Figure 4B shows the same accuracy data but as a function of viewing duration for each motion strength. Here the data combine all base rates. Note that accuracy improves as a function of viewing duration for all informative (> 0%) coherences (Eq. 19; *p* < 10^−8^, t-test, *H*_0_ : *β*_2_ = 0). This improvement is consistent with bounded accumulation of noisy evidence, where the bound curtails improvement at longer viewing durations, consistent with previous studies (Kiani et al., 2008; Kang et al., 2017). These two graphs are informative cross sections of a rich data set.

**Figure 4.**
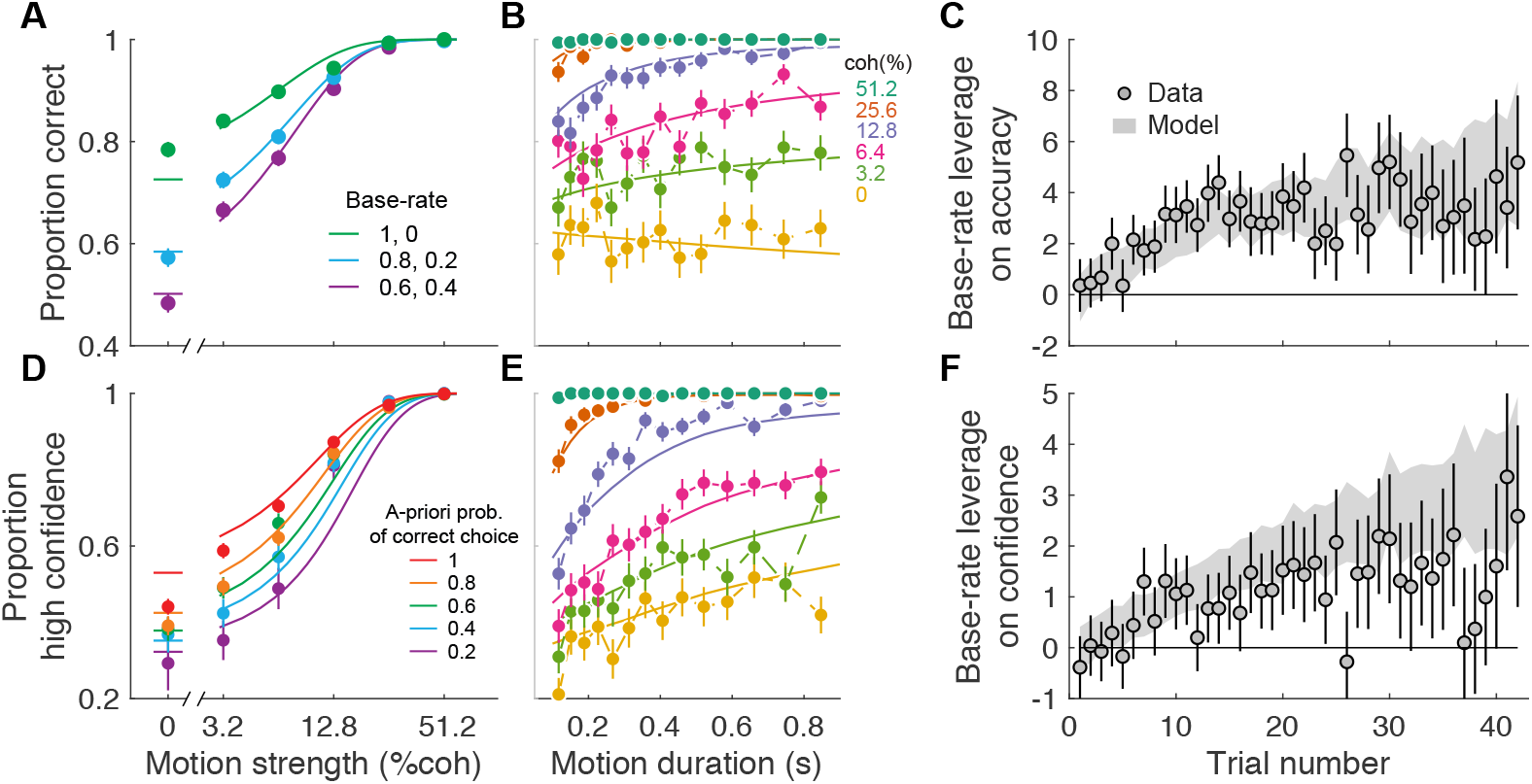
Fits of the Bayesian model to choice and confidence. (**A**) Proportion of correct responses as a function of motion strength, split by the base rate of the block (same data as Fig. 2B). Solid lines are model fits. (**B**) Choice accuracy as a function of duration of the motion stimulus, split by motion strength. Symbols are mean ±s.e., and solid lines are model fits. Points are quantiles (~157 trials per point). (**C**) The influence of the base rate on accuracy increased with trial number within blocks. Symbols are the fitted coefficients (±s.e.) from logistic regression fits to the data (*β*_6_, equation 19). The shaded area is the s.e. estimated from the model (see Methods). (**D**) Proportion of high-confidence responses on correct trials as a function of motion and base rate. The colors represent different levels of bias relative to the chosen direction (similar to Figure 2B). Symbols are mean ±s.e. across trials. Solid lines are model fits. (Figure S3 shows the error trials.) (**E**) Proportion of high-confidence responses as a function of the duration of the motion stimulus. Correct trials only. Same grouping of trials as in panel B. (**F**) The influence of the base rate on proportion of high-confidence responses increased with trial number within blocks. Symbols are the fitted coefficients (±s.e.) from logistic regression fits to the data (*β*_9_, equation 20). The shaded area is the s.e. estimated from the model (see Methods).

Knowledge of the base rate, *B*, was acquired gradually during each block of trials. Figure 4C demonstrates the time course of the changes in choice accuracy. We used logistic regression to estimate the leverage of base rate on accuracy for each trial in a block (Methods, Eq. 19). The ordinate shows the leverage of *B* on accuracy after accounting for motion strength. It supports the model free assertion that bias-dependent accuracy is learned over the course of the block (Eq. 19; *p* < 10^−8^, likelihood-ratio test, *H*_0_: all *β*_6_ = 0), and it shows that this rate is consistent with expectations of the Bayesian model (gray shading).

The model was also fit to explain the subject’s confidence in the direction report. For each subject, we tried to explain the probability that their confidence was high or low, relative to a criterion setting (see Methods). The model produces an estimate of confidence that depends on the stimulus (strength and duration), the subject’s choice, and the model’s current estimate of the base rate (Figure 3C). It did not incorporate the subject’s belief reports. The lower row of Figure 4 shows the confidence fits in a way that parallels the accuracy analyses. The trial groupings are the same as in the corresponding accuracy plots (upper row), with one exception. The block base rate (panel D) are expressed as the *a priori* probability that the direction of motion supports the choice that the participant made (correct choices only), as in Figure 2C. The model captures the important features of the data. Larger *a priori* probability of a correct choice increased confidence, and this effect was more apparent at the weaker motion strengths (Fig. 4D). Confidence also varies as a function of stimulus motion strength and duration (Fig. 4E). There is some mismatch with the model at 12.8% coherence (Fig. 4D) especially for the 300-600 ms durations (Fig. 4E). The effects build up gradually as a function of trial number in the block (Fig. 4F), showing that this rate is consistent with expectations of the Bayesian model (gray shading).

We compared the Bayesian model against two alternatives, which differ in the way knowledge of the base rates is updated across trials. As indicated in Figure 3C, a Bayesian observer will update her knowledge about the base rate after each trial using the expression,

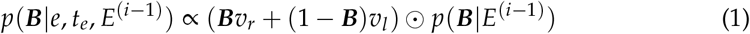

where *v_r_* = 1 − *v_l_* is equal to the counterfactual posterior for rightward motion, *p_u_*(*R*|*e, t_e_*). The symbol ⨀ indicates pointwise multiplication. In the first alternative, the participant updates *p*(*B*) based on the frequency of left and right choices, weighting all choices equally (i.e., ignoring confidence). This is implemented by making the evidence *v_r_* equal to 1 for right choices and 0 for left. We refer to this model as the Choice-only model. In the second alternative model, the evidence for updating the bias is the confidence that the participant reports in each trial, rather than the counterfactual confidence that would have been reported under a neutral prior. To model this we made *v_r_* equal to the confidence that the subjects report when they chose rightward, thus equal to *p*(*R*|*e, t_e_, E*^(*i*−1)^), and to one minus confidence when subjects chose left. We refer to this model as the Reported-confidence model. As mentioned earlier, this model is suboptimal because the evidence for updating the bias is corrupted by the bias itself.

A model comparison (Table 1) showed that the Bayesian model is the one that best fit the data for all participants. The three models have the same number of parameters (Table 2) and thus the same result is obtained with measures that penalize the goodness-of-fit by the number of free parameters (e.g., the BIC or AIC). The model comparison supports the interpretation that participants used a graded measure of certainty in the decision to update their beliefs about block bias. The support for the Bayesian model is particularly strong when we combine the likelihoods over the different participants. Note that the support for the Bayesian model derives from a model comparison which uses only the choices and the confidence in the motion direction decision.

**Table 1.** Differences in log-likelihood relative to best model for each subject. The best model for each subject is highlighted in bold.

|  | Participants |  |  |  |
| --- | --- | --- | --- | --- |
|  | S1 | S2 | S3 | Combined |
| Choice-only model | 36 | 8 | 4 | 48 |
| Reported-confidence model | 42 | 7 | 8 | 57 |
| Bayesian model | **0** | **0** | **0** | **0** |

**Table 2.** Best-fit parameters for the Bayesian model.

|  |  | Participants |  |  |
| --- | --- | --- | --- | --- |
|  |  | S1 | S2 | S3 |
| $\kappa$ | signal-to-noise | 19.64 | 23.29 | 18.01 |
| $A$ | bound height | 1.47 | 1.08 | 0.75 |
| $\omega_2$ | weight for 2nd and 5th elements of $p_0(B)$ | 0.04 | 4.9 | 0.3 |
| $\omega_3$ | weight for 1st and 6th elements of $p_0(B)$ | 0.03 | 0.41 | 0.01 |
| shrink | compression toward center of scale | 0 | 0.45 | 0.28 |
| $\phi$ | high/low confidence separatrix | 0.72 | 0.77 | 0.74 |

### The evidence for belief updating: an empirical approach

The three models represent distinct alternatives for updating: Bayesian, Choice-only, and Reported-confidence (i.e. counts weighted by confidence). The model comparison based on fits to choice and confidence provide support for true Bayesian, but the exercise fails to capture the qualitative differences in these models. Here we pursue a more general approach (Figure 5). Each of the three alternatives are instantiated by the term *v_d_* in equation 1 (where *d* stands for the chosen direction; i.e. *v_d_* = *v_r_* or *v_l_*) to update the prior over the possible base rates, *p*(*B*). These can be viewed as a mapping from two quantities to *v_d_*. These quantities are (*i*) the confidence that participants report in each trial, *p*(*d*|*e, t_e_, E*^(*i*−1)^), and (*ii*) the prior expectation that the subject had about the direction of motion before seeing the actual stimulus presented on the trial, *p*(*d*|*E*^(*i*−1)^). These quantities are the abscissa and ordinate of the graphs in Figure 5A-D. Panel A shows the map of *V_d_* in the Bayesian model, which uses the counterfactual posterior. Panels B and C show the mappings for the two alternatives. In the Reported-confidence model (Fig. 5B), the evidence *v_d_* is simply the reported confidence and does not depend on the prior expectation about motion direction. In the Ignore-confidence model, the evidence for updating the beliefs is the direction of the choice and therefore constant and independent of both confidence and prior expectation about motion direction (Figure 5C). These maps in panels A-C are not fits to data, but illustrations of the term that is used to update *p*(*B*) in each of the three models.

**Figure 5.**
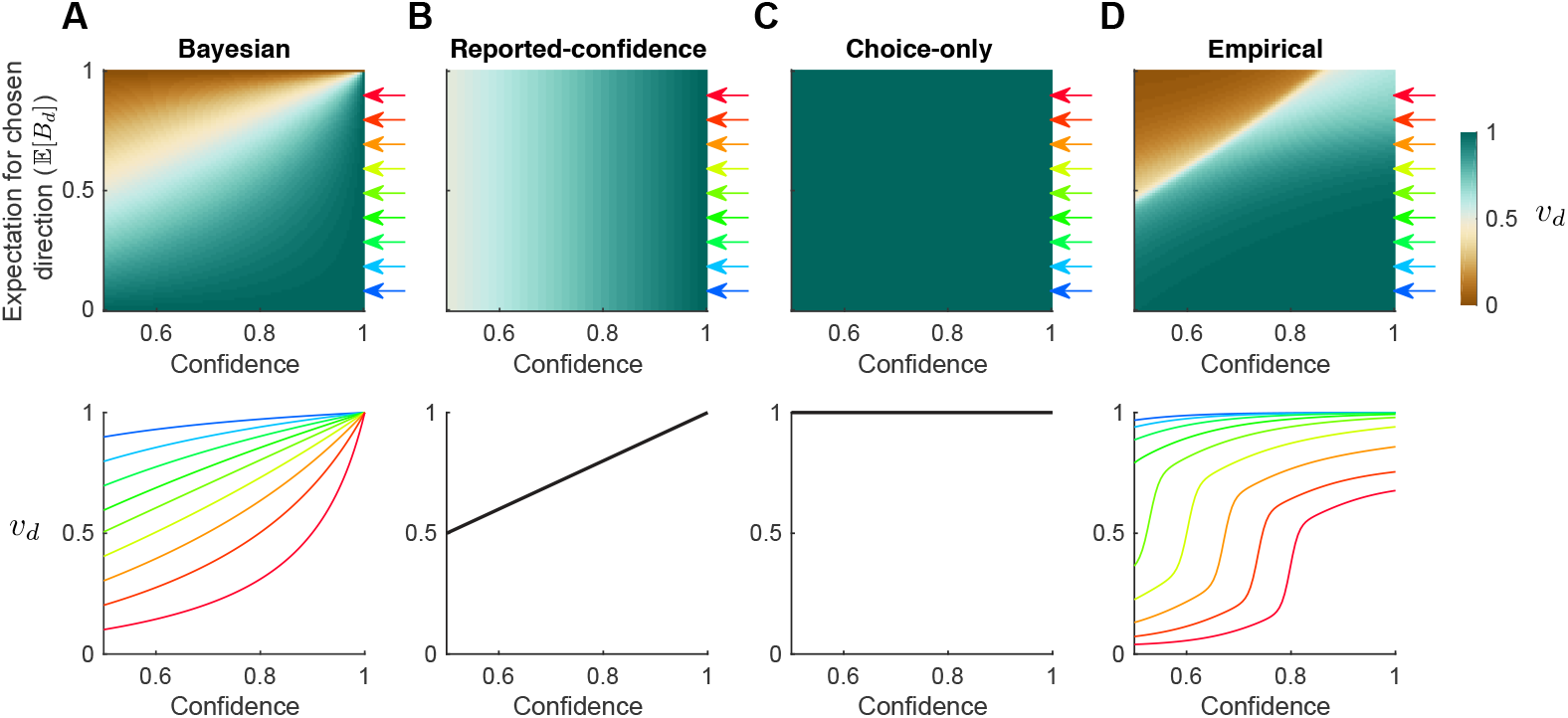
Descriptive rendering of the rule to update knowledge of the base rate. The Bayesian model and its alternatives can be characterized by the way they update *p*(*B*) based on the choice-confidence and the expectation of the base rate, expressed relative to the choice taken 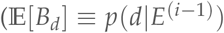, similar to a priori probability correct). The strength of the update is parameterized by *v_d_*. The top row shows the value of *v_d_* for each possible combination of confidence and expectation, for the different models. The bottom row shows the same data but only for 9 different values of the expectation over direction, which are indicated by the colored arrows (0.1 to 0.9 in 0.1 intervals). (**A**) For the Bayesian model, *v_d_* is the counterfactual confidence which is a function of both reported confidence and *p*(*d*|*E*^(*i*−1)^). (**B**) For the Reported-confidence model, *v_d_* only depends on the reported confidence (hence all the curves in the lower panel are superimposed on the identity line). (**C**) For the Choice-only model, *v_d_* is always equal to 1. (**D**) For the empirical model, we used a logistic function to specify a sigmoidal mapping of reported confidence and *p*(*d*|*E*^(*i*−1^)) to *v_d_* that best fit the data. Like the Bayesian model, confidence and *p*(*d*|*E*^(*i*−1)^) jointly determine the value of *v_d_*.

In Figure 5D, we reverse engineer this process by searching for mappings that best fit the data. We used a logistic function of the two quantities and their interaction to make new mappings of *v_d_* and used them to fit the data. The four parameters of the logistic (Eq. 16) were incorporated into the same fitting procedure used earlier, which attempts to account for each subject’s choice and confidence, which we term an empirical map. It resembles the Bayesian map in panel A. Note that for both maps, the evidence used to update the bias is stronger when the choice is made with higher confidence (Figure 5A&D). For a fixed level of reported confidence, the counterfactual confidence is also greater when the choice opposes the subject’s prior expectation about the direction of motion. Intuitively, this is because the reported confidence is a function of both the stimulus strength and the prior knowledge. Therefor, if the prior opposes the choice (expectation < 0.5), then the confidence that the subject would have under a neutral prior (expectation = 0.5) must be greater than the one reported.

Having provided further support for the Bayesian model (counterfactual confidence) we now use this model to predict the subjects’ belief reports.

### The Bayesian model predicts the belief reports

To establish fits of the Bayesian model on the basis of each subject’s choices and confidence, we relied on an estimate of the base rate (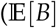, Fig. 3B), derived from the distribution, *p*(*B*). This distribution was updated in accordance with Bayesian theory using the previous decision and counterfactual confidence. However, subjects reported their belief that the block was biased to the right or left. We did not use these reports to fit the model. We can now ask how well the model predicted these reports.

In the model, the belief is obtained by summing *p*(*B*) for values of *B* greater than 0.5 (Fig. 3C),

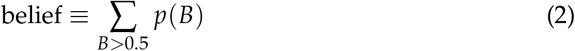

Figure 6A shows the predictions and data for the belief that the block has a rightward bias, for a few individual blocks. Each panel shows data from one randomly chosen block (colored trace). The grey lines show the belief produced from simulations of the block, using the same stimuli in the same order of the block. The simulated trials lead to different states of the accumulated noisy evidence, hence different choices and confidence, which in turn affect the evolution of *p*(*B*). Each grey line shows the evolution of belief for a different run of the model. For both the model and the participants, the belief after the first decision is close to ½. As more decisions are made, the belief tends to become more certain.

**Figure 6.**
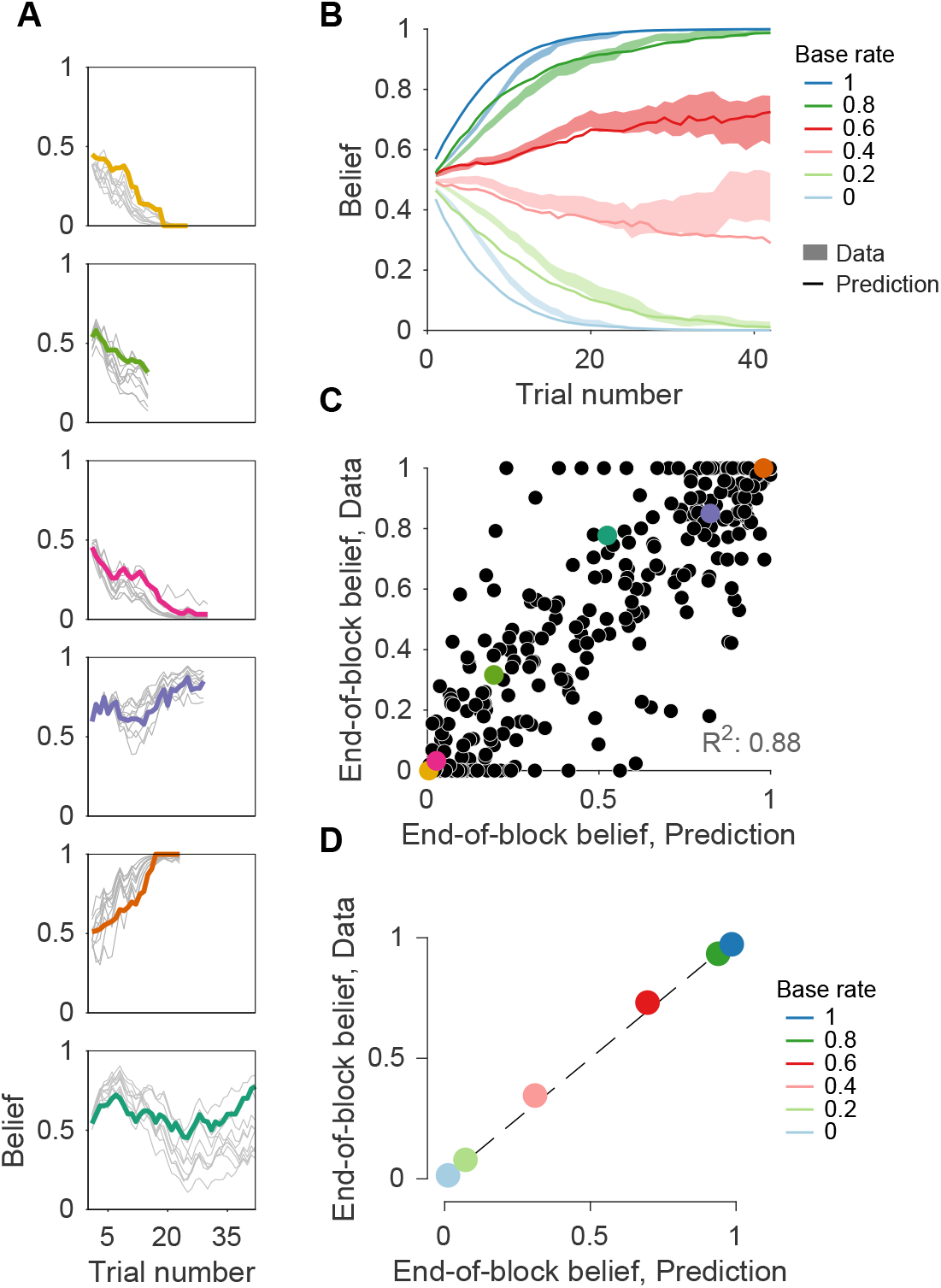
The Bayesian model predicts belief reports. (**A**) Evolution of beliefs for 6 example blocks. The colored thick lines are the data. The thin lines represent predictions from 10 simulations of the Bayesian model using the same sequence of trials (motion strength, duration and direction) as seen by the subjects. They differ because the evidence is sampled randomly on each trial ({*e, t_e_*}), which may lead to differences in choice and confidence. (**B**) Evolution of belief as a function of the trial number within the block. Lines are predictions of the model, and the shaded areas show the standard error of the data (as in Figure 2D). (**C**) Predicted belief for the last trial of each block plotted against the data from the same block. The colored circles identify the six blocks shown in panel A. (D) Same comparison as in C but averaged by actual base rate. Error bars are s.e. (usually smaller than the data points); dashed line is identity.

This is seen more clearly when we average the belief for the six different levels of base rate (Figure 2C), redrawn here as colored areas showing the s.e.m. across trials (Fig. 6B). The thin lines in Figure 6B are the predicted time course from the Bayesian model. Although subjects were slower to incorporate the bias than predicted, by trial 20 the data and predictions are almost perfectly aligned. This agreement was evident across the individual blocks. To quantify this, we compared the reported belief on the last trial of each block to the predicted belief on these trials (e.g., the average of the termination of the model simulation in panel A). The scatter plot (Fig. 6C) displays this comparison for all 485 blocks (the examples in A are identified by color; *R*^2^ = 0.88). Moreover, the average belief on the last trial of each block (data points in Fig. 6D) was almost identical to the predictions of the Bayesian model when grouped by base rate. The agreement between data and predictions in Figure 6 is remarkable, considering that the predictions were based only on the fits of the model to the direction choice and confidence.

Another prediction of the Bayesian model is that stronger motion (higher coherence and longer stimuli), which is associated with higher confidence on average, should lead to a larger change in the belief that the block is biased to the right or left. Figure 7A-B corroborates this prediction. For each coherence and for the two choices, we show the change in belief relative to the current belief reported at the end of the previous trial. Not surprisingly, the belief tends to increase when subjects choose rightward motion, and decrease when they choose leftward motion. Further, because the scale is bounded, the changes approach zero on the extremes of the scale. More interestingly, for the same level of the current belief, stronger motion leads to larger changes in belief (Eq. 21; *p* < 10^−8^, t-test, *H*_0_ : *β*_1_ = 0). Similarly, splitting trials by the median duration of the motion stimulus and grouping all coherences shows that longer durations are associated with larger changes in belief (Eq. 21; *p* < 10^−8^, t-test, *H*_0_ : *β*_2_ = 0; Fig. 7B). The model predicts a similar pattern (Fig. 7C-D) although the predicted changes are larger than what is seen in the data. One factor that contributes to this mismatch is that participants do not indicate any change of belief on approximately 30% of trials. This could reflect a resistance to make or report small changes in belief. Importantly, and regardless of the difference in absolute magnitude, the relationship between motion strength and change in belief in Figure 6 is incompatible with models in which only choice—and not the certainty of the choice—informs the revision of belief.

**Figure 7.**
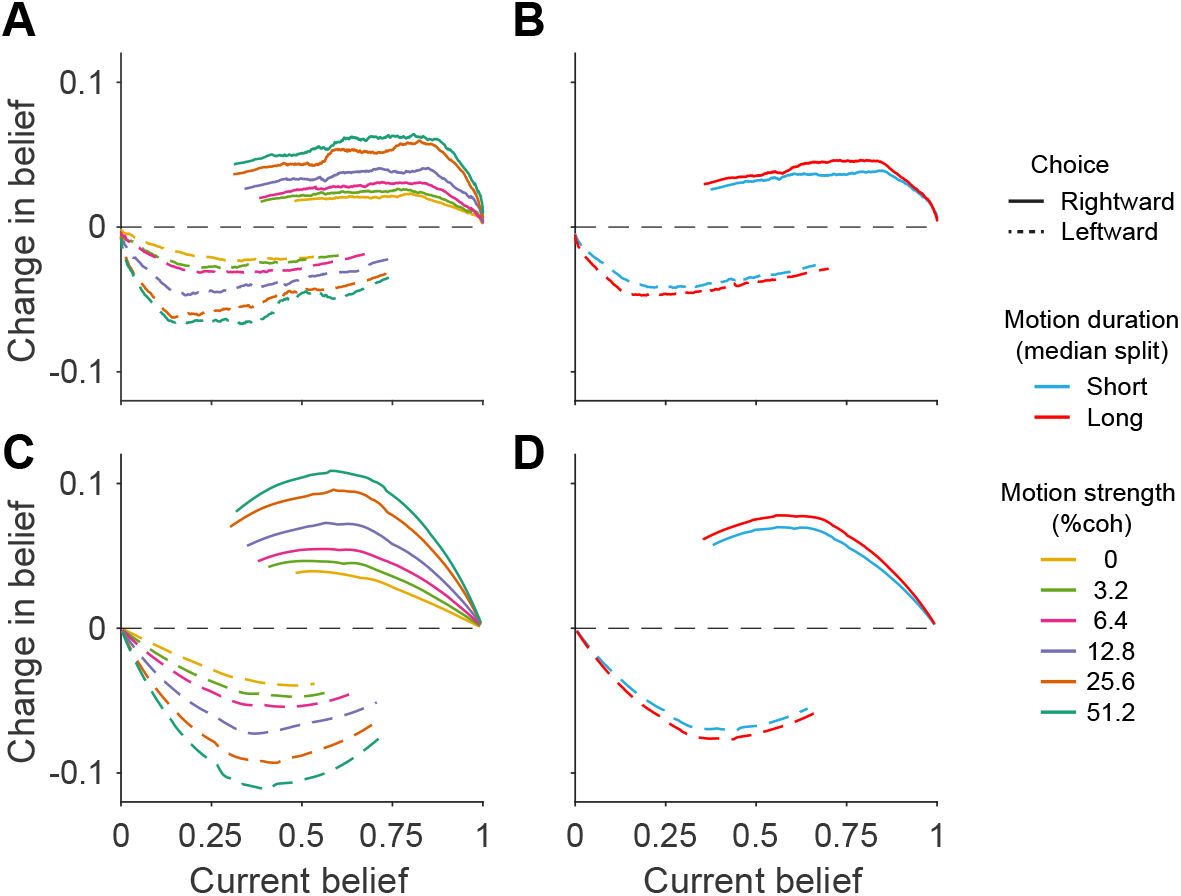
Change in belief depends on current belief, choice, motion strength and duration. All panels depict the average change in belief from one trial to the next, as a function of the current belief (correct trials only). (**A**) Data split by choice (solid vs. dashed line) and motion strength (color). Data are a running means of N = 200 trials. (**B**) Data split by choice and duration (color). (**C & D**) Same as A & B, for model simulations.

An assumption of the model is that subjects represent the base rate as a distribution *p*(*B*) which they update on each trial and from which they calculate the belief that they report on that trial. The belief that the block is biased in favor of right does not imply that it is biased strongly or weakly. For example, a decision maker may report a belief of 1 (i.e. certain the bias is rightward), but we do not know, from this observation, if she considers the base rate to be 0.6 or 1. Moreover, the same belief could be associated with different shapes of *p*(*B*) and different expectations of *B*.

We know from the choice and confidence reports that subjects behave as if they represent different expectations of *B*. To determine if participants represent different distributions of base rate even when they report the same degree of belief, we analyzed the choices on 0% coherence trials in which they were close to certain about the bias of the block (> 0.95 or < 0.05 on the previous trial). If the subject’s representation of base rate matches that of the block, then on these trials with little net motion, the subject should report more frequently in the direction of bias when the base rate is more informative, despite the belief being equally strong. Figure 8 supports this prediction. Subjects were more likely to report the direction of the belief when the base rate was more extreme (Figure 8; Eq. 22; *p* < 10^−6^, t-test, *H*_0_ : *β*_1_ = 0). A similar trend was observed in simulations of the Bayesian model. This analysis presents model-free evidence that participants represent not only the quantity that they are asked to report (belief) but also a measure of the magnitude of the base rate.

**Figure 8.**
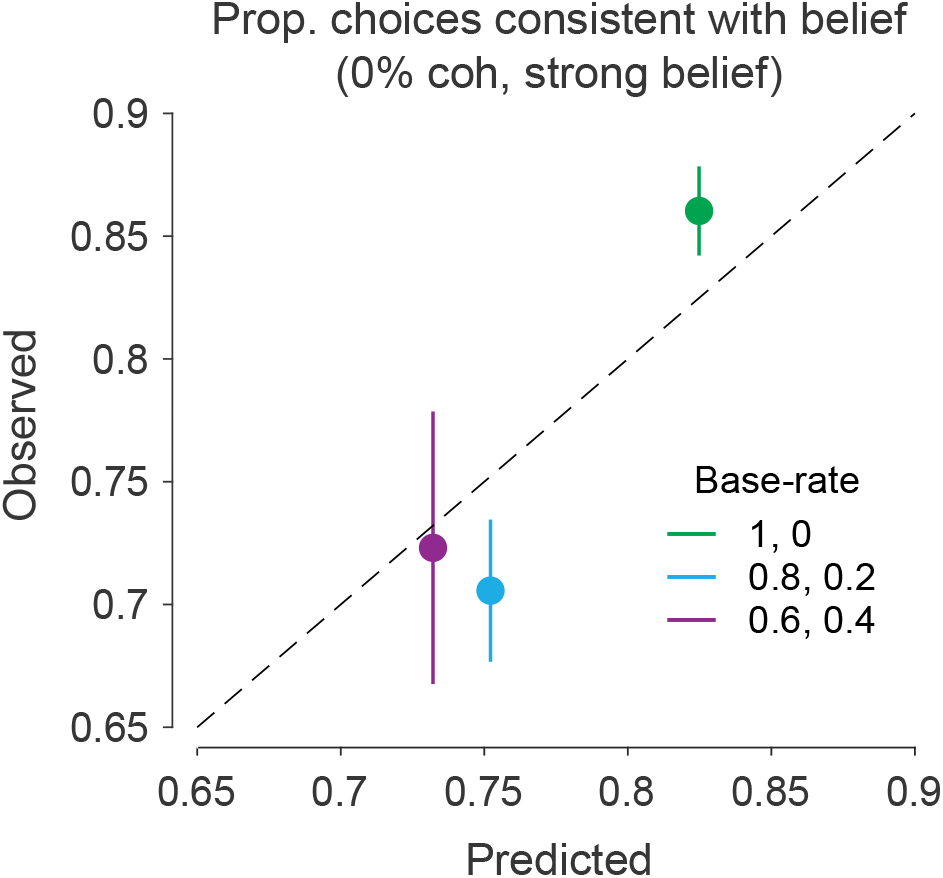
Belief is not a report of the base rate. The graph depicts only trials in which the subjects were more than 95% certain that the block was biased. Scatter plot compares model predictions and observed choice frequencies for the three levels of informativeness of the base rate (color). The choices made on 0% coherence motion should be strongly influenced by their estimate of base rate, but the choice bias is modest for less informative base rates. (N = 65, 248, 365 trials for base rates of low, intermediate and high informativeness; predictions based on 200 simulations). Error bars are s.e.m.

Our Bayesian model incorporates one element of sub-optimality. Although subjects were informed that the distribution over base rates was discrete and uniform, we allowed for the possibility that the prior assumed by the subjects was not veridical. We approximated this initial prior, *p*_0_(*B*), with 3 parameters, which were estimated in the fits to the direction choice and confidence (see Methods). All three subjects appear to have assigned a greater initial probability to base rates away from the extremes (Figure S4). This might imply that subjects were unable to override a predisposition that sequences of random samples tend to be only weakly biased, on average—an arguably sensible prior over prior probability distributions—despite direct verbal instruction. The deviation from optimality was only marginally detrimental to performance. For example, the proportion of correct responses was 85.01% in the data and 85.2% in simulations of the Bayesian model, whereas we estimate that the uniform prior would yield 87.3% correct.

## Discussion

Base rate, prevalence and prior probability distribution are examples of statistical regularities that ought to shape our decisions. A decision maker must learn these regularities from sources, such as census or epidemiological research, or through learning. We have shown that decision makers can infer such regularities from their own decisions, and use the evolving knowledge of the base rate—its prior probability distribution—to make better decisions, even during the learning process itself. Remarkably, they did this without feedback about the validity of their decisions, which were for the most part difficult. Indeed over half of the decisions were about weak motion that would under neutral priors give rise to correct choices on less than 75% of trials, that is, just better than guessing. Such difficult trials are the very ones that benefit the most from knowledge of the base rate. We showed that subjects incorporated an evolving estimate of base rate into these decisions and their confidence, while at the same time exploiting the decision to update knowledge of the base rate.

We observed that the influence of the base rate on our three behavioral measures—decision, confidence and belief—developed gradually during the block. This was evident in the two ways by which we evaluated the learning of the base rates: implicitly, through their influence on decision and confidence; and explicitly, through the belief report. Both methods gave consistent results. Indeed, we were able to fit a model to the decision and confidence reports, and predict how the belief evolved during the block. This suggests that the influence of bias on the implicit and explicit reports is mediated by a common estimate, which in our model is represented by a probability distribution of possible base rates that is updated from one decision to the next. It also shows that people have introspective access to quantities that are used in the computations of a hierarchical probabilistic model, and therefore subjective reports can be used to constrain models of decision making (Kang et al., 2017).

A theoretical analysis showed that the evidence for the revision of beliefs is a ‘counter-factual’ form of confidence, understood as the confidence that the subjects would have if left and right were equally likely (i.e., a neutral prior), and not the confidence that the subjects report. The intuition for this distinction is simple: the evidence used to modify an hypothesis should not be altered by the decision maker’s previous belief about the veracity of the hypothesis. A comparison of models supported the conclusion that subjects indeed used the counterfactual confidence to update their bias. This conclusion was confirmed with a separate model in which we directly search over the space of transformations that map confidence and bias strength on the evidence that is used to update their knowledge of the base rate (Fig. 5). The use of a counterfactual confidence for updating the bias implies the existence of different probabilistic quantities associated with the same state of accumulated evidence, a distinction that may be related to the one between confidence and visibility that has been recently reported (Rausch & Zehetleitner, 2016).

The concept of a counterfactual posterior, and the related counterfactual confidence, demands some justification. Mathematically the counterfactual posterior is simply the likelihood. Moreover, just because we factor the update rule to highlight this quantity does not automatically imply the the brain uses it. We believe the interpretation has merit, however, because the counterfactual posterior can be approximated by a logistic function of evidence and time (Kiani & Shadlen, 2009; Drugowitsch et al., 2012; Shadlen et al., 2006). Indeed it is obvious that a representation of counterfactual confidence is essential in many types of real world decisions. For example, if instead of manipulating the prior probability of right and left, we had altered the reward such that all correct rightward choices were rewarded more than leftward, it would induce a rightward bias. In this circumstance, it would be sensible to express both a confidence that we made the better choice and a confidence that the motion was to the right. For example when confronted with weak motion, we would be confident that a rightward choice was best while simultaneously less confident that the net direction was to the right. The dual representation of confidence is intuitive when value induces a bias but less so when prior probability (base rate) does so, because the prior is about the motion itself. What we show here is that such a dual representation of confidence in our decision and counterfactual confidence allows us to both respond as accurately as possible and learn about the statistical regularities of the environment.

Our task represents an attempt to extend our understanding of the neurobiology of simple decisions to tasks that have hierarchical structure. One tempting idea is that the same process of evidence accumulation that allows one to make a simple perceptual decision extends to higher levels and longer time scales (Kim et al., 2017; Purcell & Kiani, 2016; Glaze et al., 2015). In the abstract this is true because both are effectively Bayesian, but this insight deserves further scrutiny. On the one hand, the decision variable (DV) on a single trial is the accumulation of evidence for rightward, say, and against leftward. The accumulation defines a random walk such that at any moment the DV is proportional to the log odds that rightward is correct, where the constant of proportionality is a straightforward function of time (Drugowitsch et al., 2012). To incorporate bias, it is possible to add the log of the prior odds, scaled as a function of time, and there is support for such a mechanism at the level of single neuron recordings (Hanks et al., 2011).

On the other hand, to update *p*(*B*) in a similar manner would imply the accumulation of a scalar quantity throughout multiple trials to determine block bias. This strategy would be optimal if the decision maker were to receive unambiguous feedback about the true direction on each trial (Laplace, 1814). However, when the decisions vary in difficulty (and thus in confidence) there is no scalar quantity that the subjects can accumulate from one trial to the next that preserves all information about *p*(*B*). To give an intuition, suppose the subject used counterfactual confidence, signed by choice, to update *p*(*B*), and consider two scenarios. (1) A decision maker chooses rightward on the first trial and leftward on the second, both decisions made with full confidence. (2) The same choices ensue but with very low levels of confidence. In both scenarios, the running tally would be zero. However, in the first scenario the subject reduced her uncertainty more than in the second scenario, since she learned that the block has both right stimuli and left stimuli and therefore can assign low probability to the most extreme biases. This knowledge is captured by the normative propagation of *p*(*B*) but not by any scalar accumulation strategy. However, In reality the DV for single decisions is represented as a race between two accumulation processes—more if the decision is between 3 or more choices—and the distribution of *p*(*B*) may be approximated with fewer parameters (e.g., beta distribution).

The present study complements several recent studies which have addressed the manner in which a decision maker can sense a change in the statistical structure of the environment (Summerfield et al., 2011; Behrens et al., 2007; Norton et al., 2017; Nassar et al., 2010; Yu & Cohen, 2008; Anderson & Carpenter, 2006), similar to the change in base rate that occurs (with probability 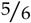) at the beginning of each block in our experiment. Whereas we provided explicit instruction that the block changed and no feedback about the decisions made during the block, these complementary studies provide no explicit feedback that the environment has changed. Instead they provide feedback about the success or failure of the decisions made in the new environment (cf. (Kim et al., 2017)). The common theme is that statistical inference occurs over two temporal scales, one concerning individual decisions and the other the statistical regularities about the environment in which these decisions are undertaken. The key contribution of the present study is to show that different forms of confidence apply to the two time scales: confidence that the decision is correct (which incorporates knowledge of base rate) and counterfactual confidence in the direction to update the prior over base rate, and confidence in the bias of the block (i.e. belief). In the tomato problem introduced earlier, the sorter would apply whatever knowledge of the base rate she has to each tomato. It would tend to influence the borderline cases most. With each decision, knowledge of the proportion of ripe and unripe would be updated using confidence in the tomato itself, stripped as it were, from the influence of base rate, as if imagining a box with equal proportions of ripe and unripe fruit. The sorter might use this evolving knowledge to tell a co-worker that a crate seems to contain mostly unripe fruit and the degree to which she believes this to be true. She could also report her estimate of the proportion itself, but as in our experiment, that will also be evident in the decision about the next sample and the confidence that it has been sorted correctly.

Without immediate feedback from the world, confidence is all we know about the veridicality of our assertions. Recent progress in understanding the mechanisms for deciding and assigning a degree of confidence to isolated decisions has set the stage to study the role of confidence in more complex tasks, especially those that involve multiple steps (Gold & Stocker, 2017; Zylberberg et al., 2011). It has been suggested that confidence plays a key role in assigning blame to different sources of evidence after an error (Purcell & Kiani, 2016; McGuire et al., 2014), controlling how much effort and time to invest in a decision that depends on the success of a previous decision (van den Berg et al., 2016), combining decisions in a hierarchy to maximize reward (Lorteije et al., 2015), and guiding perceptual learning in the absence of feedback (Guggenmos et al., 2016). Confidence may also play a role in combining individual opinions with those of a group according to their reliabilities (De Martino et al., 2017; Park et al., 2017; Bahrami et al., 2010). Our work adds to these studies by showing that confidence mediates the bidirectional process by which decisions both inform and are informed by a dynamic estimate of a base rate. It does so by drawing implicitly on counterfactual knowledge about confidence associated with a setting that is inapplicable to the decision at hand. Counterfactual reasoning is thought to play a role in slow, deliberative decision making, where it relies on narrative devices or playing through imagined scenarios. It is intriguing to consider that a more automatic form of counterfactual reasoning, such as we have demonstrated here, arises in a “thinking fast” (Kahneman, 2011) mode, where it conforms to normative Bayesian principles.

## Methods

### Behavioral Task

Three human participants (2 male, 1 female; age 24-35) gave informed consent and participated in the experiment. The experimental protocol was approved by the local ethics committee (Institutional Review Board of Columbia University Medical Center). Stimuli were displayed on a CRT monitor (ViewSonic, PF790) with a refresh rate of 75 Hz. A head and chin rest ensured that the distance between the participants eyes and the monitor’s screen was 48 cm. We tracked the position of the eye with an Eyelink 1000 eye-tracker.

Subjects performed a motion direction discrimination task in blocks of trials. They were given no information on their performance until the end of each block. The number of trials in each block was sampled from a truncated geometric distribution 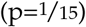 with a minimum of 15 trials and a maximum of 42 trials. Within each block of trials, one of the motion directions (leftward or rightward) was more probable than the other. The base rate of each block was sampled with equal probability from the set *B* ∈ [0, 0.2, 0.4, 0.6, 0.8, 1], where *B* indicate the proportion of rightward directions. That is, given a block of *n_tr_* trials, the number of trials with rightward motion was *B* × *n_tr_*, rounded to the nearest integer. The order of trials was randomly permuted. Subjects were informed about the set of base rates and that they were equally probable for each block, but they were not informed about the base rate of each block until the block was completed.

Each trial started with participants fixating a small point (diameter of 0.33 deg) at the center of the screen. After 0.5 s, two target arcs (2*π*/3 radians) were presented each side of the fixation point (Figure 1; similar to (Zylberberg et al., 2016)). The circular arcs each had a radius 10 deg of visual angle. Subjects had to select the left and right targets to indicate leftward and rightward motion, respectively. Subjects used the upper extreme of targets to indicate full decision certainty and the lower extreme to indicate guessing. Intermediate values represent intermediate levels of certainty. As a visual aid, the targets were colored from green to red from top to bottom (Fig. 1).

After a variable duration of 0.3 - 1s (truncated exponential; *τ* = 0.5 s), the random dot motion was presented in a virtual aperture centered at fixation and subtending 5 degrees of visual angle. The dot density was 16.7 dots/deg^2^/s and the displacement of the coherently moving dots produced an apparent speed of 5 deg/s. In each video frame, a subset of the dots presented 40 ms earlier were displaced toward the right or left target, depending on the motion direction for that trial. The details of the stimulus generation process can be found in previous studies (e.g., (Roitman & Shadlen, 2002)). The probability that a dot moved coherently as opposed to randomly varied from trial to trial according to the motion strength (|*c*|), sampled uniformly from the set [0, 3.2, 6.4, 12.8, 25.6, 51.2]%. To keep the motion strength approximately balanced within each block, we constructed a list of motion coherences by repeating the six unique coherence values until the first multiple of six that is larger than *n_tr_*; then we shuffled these motion coherences, and assign the first *n_tr_* values to the trials in the block. Note that the distribution of motion strengths within a block is independent of the base rate.

The random dot motion disappeared after 0.1-0.9 s (truncated exponential, *τ* = 0.6 s), together with the fixation point, and the participant then reported her decision with a computer mouse. Subjects were free to move the cursor until satisfied with their choice and confidence report (signaled by pressing the spacebar). As a visual aid, when the cursor approached one of the arcs, a marker was shown on the arc that was closest to the cursor position together with a number between 50 and 100, indicating the probability (as percentage). See Movie S1 for the actual stimulus display.

After the choice and confidence reports about motion direction, subjects reported the degree to which they considered the block to be biased in one direction or the other, indicated on a continuous horizontal scale. The leftmost extreme indicated full certainty in a left bias (belief = 0) and the rightmost extreme indicated full certainty in a right bias (belief = 1). The in-between points represented intermediate levels of certainty, with the center of the scale indicating maximal uncertainty, meaning that subjects considered equally likely that the block was biased in one direction or the other. We used color gradients as a visual aid, with a scale that went from white to black and then back to white. Subjects reported their decision by moving a small triangle to the desired location before pressing the spacebar to record the belief as the current position of the triangle (see Movie S1). For the first trial of a block, the triangle was rendered at the center. For all subsequent trials it reappeared at the location where it was set in the previous trial.

Subjects were only provided with feedback after a block had been completed. Feedback comprised a single screen indicating whether the block was biased to the left or to the right, the probability that a trial was biased in this direction (60, 80 or 100%), and the proportion of trials in the block that the subject reported correctly. An example feedback screen is shown in Figure 1B.

Participants performed the task across multiple sessions (9 to 14, usually 1 session/day). They usually completed 12 blocks per session. Across days, participants completed 165, 150 and 170 blocks, respectively, for a total of 4564, 4051 and 4554 trials.

Before performing the main experiment, participants were trained to discriminate the direction of random dot motion. During training, the base rate was fixed and equal to ½. Participants first performed a version of the task in which they were only required to report the motion direction (436, 432 and 936 trials per participant). During a second stage of training, they also reported their confidence, using the same numerical scale as in the main experiment (961, 288 and 384 trials per participant).

### Model

We used a bounded evidence accumulation model to fit the choices and the confidence about motion direction. In the model, these reports are based on the state of accumulated motion evidence (decision variable, *e*) at the time (*t_e_*) a bound (at ±*A*) is reached or when the stimulus ends. The decision variable evolves according to 1-dimensional process of Brownian motion plus deterministic drift. In small time steps *δt*, the state of the decision variable is updated based on the sum of terms representing a deterministic drift and stochastic diffusion. The drift term is given by *κcδt*, where *c* is the signed motion coherence and *κ* is parameter which determines the signal-to-noise ratio. The diffusion term follows a normal distribution with a standard deviation of 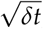 such that the variance over a 1-second stimulus is equal to 1.

The model assumes that the decision maker has access to the state of the decision variable e and the decision time *t_e_*. In the Bayesian model, the reports of choice and confidence are based on the posterior probability over motion direction, 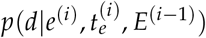, and the belief report is based on the posterior probability over base rate, 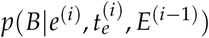. In these expressions, *d* is the motion direction, *B* is the base rate of the block, and *i* is the trial number. *E*^(*i*)^ is the tuple of all observations of *e* and *t_e_* up to trial *i* within a block: *E*^(*i*−1)^ ≡ {*e*_1..*i*−1_, *t*_*e*,1..*i*−1_}.

The posterior over the two possible directions of motion, 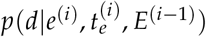, can be obtained by Bayes rule:

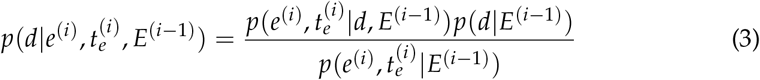

Which simplifies to:

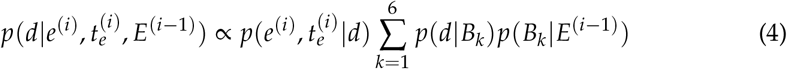

where the constant of proportionality is such that the sum of the probabilities of right and left equals 1. The sum over *k* is over the six possible values of *B*.

The likelihood in expression 4 can be computed by marginalizing over motion coherence (*c*):

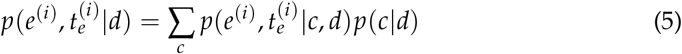

where *p*(*c*|*d*) is a uniform distribution over the coherence values that are compatible with motion direction *d*, and 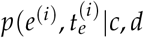 is obtained from the numerical solutions to the Fokker-Planck equation, which governs the bounded drift-diffusion (Kiani & Shadlen, 2009).

By Bayes rule, the likelihood 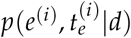 is proportional to the posterior, 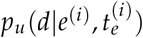, under a neutral prior. Further, *p*(*d*|*B*) is equal to *B* if *d* is rightward, and to (1 − *B*) if *d* is leftward. These substitutions lead to the expression in Figure 3B.

The choice and the confidence about motion direction are derived from the posterior 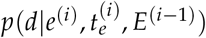. The decision maker chooses rightward if 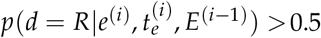 and chooses leftward otherwise.

The belief is obtained from the posterior over base rate, 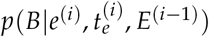. By Bayes rule:

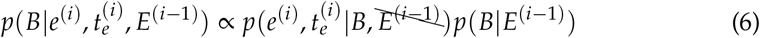

where the constant of proportionality is such that

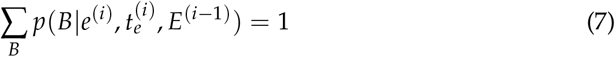

and

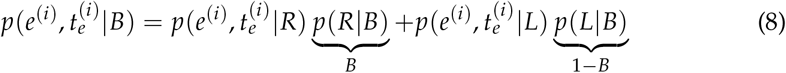

The strikethrough in equation 6 serves as a reminder that *E*^(*i*−1)^ only affects the probability of the evidence through *B*. Again, noting that 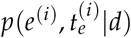 is proportional to the confidence under a neutral prior, 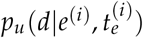, we obtain the expression in Figure 3C.

These calculations are iterated for the subsequent trials in the block, where the posterior after one trial becomes the prior for the next one:

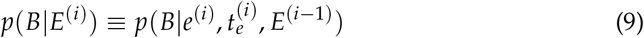

### Model fitting

The model was fit to the choice and confidence of each trial. For the first trial in a block, because the subject knows the six possible values that the bias can take and that they are equally likely, the initial distribution over bias *p*_0_(*B*) should be uniform on these discrete values. However, we assumed that *p*_0_(*B*) could be non-uniform and that the values of *B* could deviate from their true values. This was implemented with a ‘shrink’ parameter, which scaled all the values toward 0.5. If ‘shrink’ is equal to 0, the values are equal to those in the experiment. If shrink=1, all values of *B* are equal to 0.5. Note that now we are using B to refer not to the base rate of the block as defined by the experimenter, but to an internal representation of these values. We still assume that there are six possible values that *B* can take, and that they are symmetric around 0.5, and thus the full distribution is specified by two additional parameters. For fitting, the weight of the two values closer to 0.5 was clamped to 1 (before normalizing the probability distribution), and the other two weights, *ω*_2_ (for the two intermediate value of *B*) and *ω*_3_ (for the highest and lowest values of *B*), were fit to the data. The best-fitting parameters for each subject are shown in Table 2.

For fitting, we categorize the confidence reports that the subjects gave into two categories: high or low. This allowed us to treat each confidence report as a binary variable with likelihood comparable to the choice. For each subject, the lowest 30% of the confidence reports were considered low confidence. The cutoff seemed sensible given (i) the way confidence reports cluster in the scale (Fig. S2), (ii) that it approximates the proportion of low confidence reports in a similar task in which subjects categorized confidence into low or high (van den Berg et al., 2016), and (iii) that changing this value within a reasonable range does not alter the results reported here.

We performed maximum-likelihood fits of the model parameters ***θ*** separately for each subject:

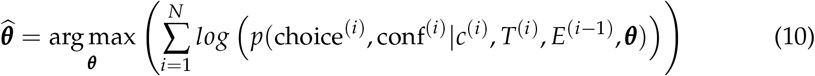

where *N* is the number of trials, choice and conf are the chosen alternative (left/right) and the category of the confidence report (high/low) on trial *i*. Note that information from the previous trials in the block affects the current one through the distribution over base rate *p*(*B*|*E*^(*i*−1)^). The joint distribution over choice and confidence is calculated by marginalizing over the values of *e* and *t_e_* that are consistent with the report. For instance, the probability of a rightward choice of low confidence is calculated as:

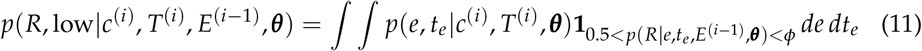

where **1**_*x*_ is an indicator function that evaluates to 1 when *x* is true and to zero otherwise, and *ϕ* is a fitted parameter—a criterion on probability correct that separates high from low confidence responses. The indicator function restrict the integration to the range of values of *e* and *t_e_* for which the probability of rightward motion is higher that 0.5 but lower than *ϕ*.

One complication in updating the probability distribution over *B* from one trial to the next is that we need to know the ratio of the likelihoods 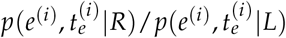, or equivalently, the ratio of the confidence in rightward and leftward choices under a neutral prior, 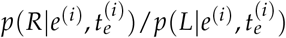. These likelihoods depend on state of accumulated evidence *e* and the decision time *t_e_* of each trial, which are not available to the experimenter. However, we can use Bayes rule to derive the ratio from the confidence reports and the distribution over base rates from the previous trial.

On trials in which subjects chose rightward, confidence is equal to 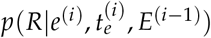 which by Bayes rule is:

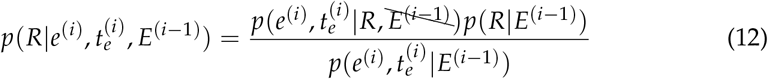

where the strikethrough makes explicit supervenience (i.e., given direction, the evidence from previous trials is irrelevant to the likelihood). Similarly,

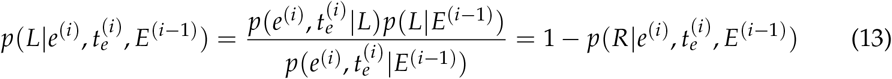

Then the ratio of the likelihoods can be derived from the confidence reported by the subject, and the prior expectation over direction of motion that is updated over trials:

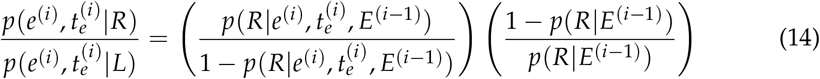

The Bayesian, Choice-only and Reported-confidence models all have the same six parameters: for the low-level motion discrimination decision, *κ, A, ϕ*; for distribution over bias: *ω*_2_, *ω*_3_ and shrink. The result of the model comparison using the log-likelihood of each model is shown in Table 1, and the best-fitting parameter values for the Bayesian model are shown in Table 2.

From the best fitting model parameters, we predicted the participants belief reports using the posterior 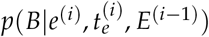. The belief that the block is biased to the right is given by

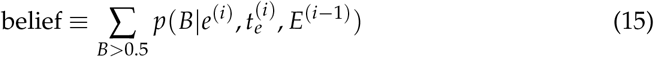

This sum is directly contrasted with the participants’ belief reports in Figure 6.

In the empirical model (Fig. 5D), we parameterized *v_d_* (from expression 1) as a logistic function with four parameters:

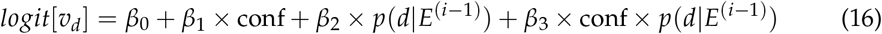

The *β* are fit simultaneously with the other 6 parameters of the model. The mapping from confidence and motion expectation to *v_d_* for the best-fitting model is shown in Figure 5D.

The model fits and predictions in Figures 4,6,7,8 were obtained from simulations of the Bayesian model using the fitted parameters. Predictions were made by running 200 simulations with the same sequence of trials as those performed by the participants. To produce the smooth lines in Figure 4A,B,D,E, we simulating each trial repeatedly with a fine grid of motion strengths (Fig. 4A,D), and stimulus durations (Fig. 4B,E). Simulations of each trial is needed because the distribution over base rates varies from trial to trial, and thus each trial is in a sense unique. As indicated in the legend, in Figure 7 the moving averages for the data are calculated over 200 trials. For the simulations, the moving averages for the data are calculated over 200 × *n_sims_* trials, after grouping trials from *n_sims_* = 200 simulations of each block.

### Data analysis

We used different linear and logistic regression models to evaluate the influence of the base rate, motion strength and viewing duration, on choice, confidence and belief. Unless otherwise noted, we used t-tests to evaluate the null hypothesis that a single regression coefficient is zero (using the parameter estimates and its standard error), and likelihood ratio tests for nested models when testing more than one regression coefficient simultaneously.

The logistic regression model used to examine the effect of base rate on choice (solid lines of Figure 2A) is:

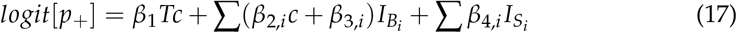

where *p*_+_ is the probability of a rightward choice, *c* is the motion coherence and *T* is the stimulus duration. The subscripted indicator variables *I* are over base rate *B* and subject *S*. For instance, *I_B_i__* is equal to 1 for trials from blocks with base rate *B_i_*, and zero otherwise. The second subscripts of the *β* parameters indicate separate parameters fit over the indexed categorical variables. The summations are all over the index *i*. The inset in Figure 2A shows the *β*_3,*j*_ coefficients and associated standard errors obtained from the logistic fits using combined data from all subjects.

To examine the influence of base rate and motion strength on confidence, we used the following linear regression model, fit using correct decisions only:

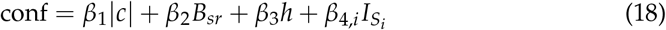

where conf is the confidence report (0.5-1 scale), *B_sr_* is the base rate relative to the choice (*B* for rightward and 1 − *B* for leftward choices), and *h* is the choice (left or right) to account for potential differences in confidence due to chosen side.

The regression model used to evaluate how accuracy depends on motion strength, motion viewing duration, trial number and base rate (Figure 4C) is:

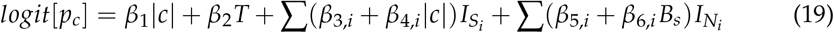

where *p_c_* is the probability of a correct response, *N* is the trial number, and *B_s_* is the informativeness of the base rate (i.e., either 0.6, 0.8, or 1). Figure 4C (top) shows the *β*_6_ coefficients.

A similar logistic regression model was fit to confidence (Figure 4F):

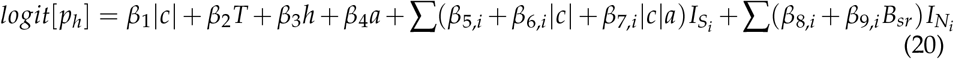

where *p_h_* is the probability of a high-confidence report, *h* is the choice (left or right) to account for potential differences in confidence due to chosen side, and *a* is the accuracy of the response (0 or 1 for incorrect and correct decisions respectively). Figure 4F (bottom) shows the *β*_9_ coefficients. The extra terms are included as potential confounders.

The same regressions were used for both data and model in Figure 4C&F. For the model, we conducted the logistic regressions independently for 200 simulations the model, and then averaged the regression coefficients. The standard errors (gray shades) were calculated as the standard deviation of the coefficients over the 200 simulations. The model was evaluated on the same sequence of trials as the participants (same direction of motion, coherence, viewing duration and order of trials).

To test the influence of motion strength on ΔBelief (Fig. 7), we fit the following linear regression model:

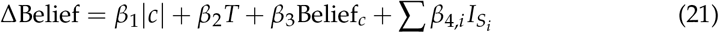

where ΔBelief is the difference between the belief reported in the current trial and the belief reported in the previous trial (or the difference from ½for the first trial of the block). To simplify the regression model, ΔBelief was multiplied by −1 when the subject chose the leftward direction of motion, and thus a positive ΔBelief indicates a change in the direction of the choice. Accordingly, Belief_*c*_ is the belief in the current trial (see Fig. 6) relative to the actual choice (i.e., Belief_*c*_ is equal to the reported belief for rightward choices, and to one minus the reported belief for leftward choices). As in Fig. 7, only correct trials were used for the regression model.

We used the following logistic regression model to test the influence of the block’s base rate on the choices for 0% coherence trials (Fig. 8):

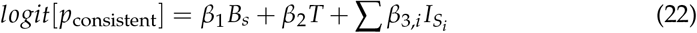

where *p*_consistent_ is the probability that the choice is made in the direction considered most likely by the subject according to the belief reported on the previous trial. As in Figure 8, in the regression analysis we only included trials of 0% motion strength and those in which the belief in the previous trial was either lower than 0.05 or higher than 0.95.

## Acknowledgments

The research was supported by the Howard Hughes Medical Institute and National Eye Institute grant R01 EY11378 to M.N.S., the Human Frontier Science Program to D.M.W. and M.N.S., the Wellcome Trust and Royal Society (Noreen Murray Professorship in Neurobiology) to D.M.W.

## Author Contributions

A.Z. designed the experiment, collected and analyzed the data and performed the computational modeling. D.M.W. and M.N.S. helped in data analysis and provided intellectual support throughout the study. A.Z., D.M.W. and M.N.S. wrote the manuscript.

## Supporting Information

**Figure S1.**
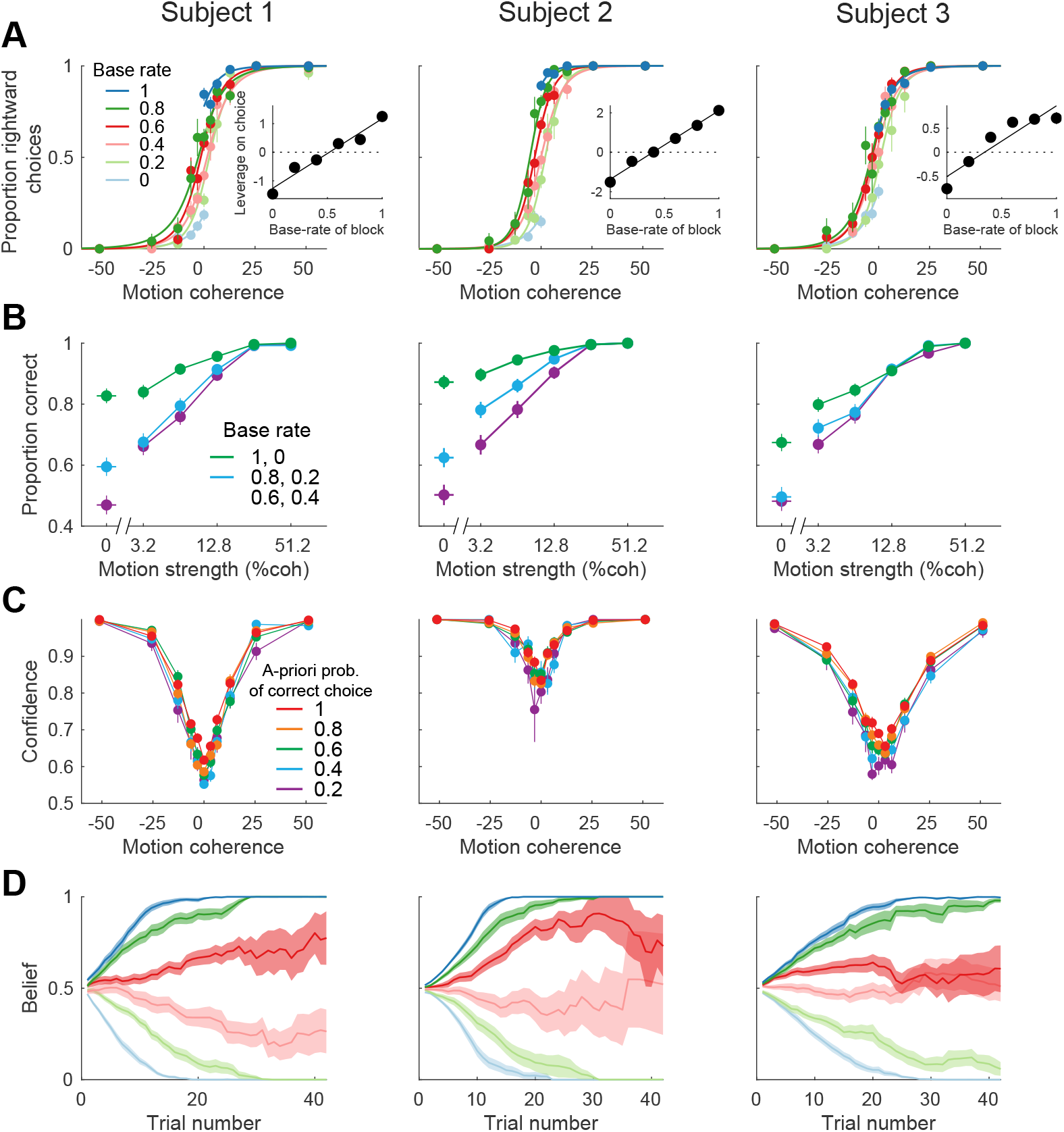
Behavioral influence of the block’s base rate. As Figure 2, but shown separately for each participant (columns). Rows correspond to panels A-D of Figure 2.

**Figure S2.**
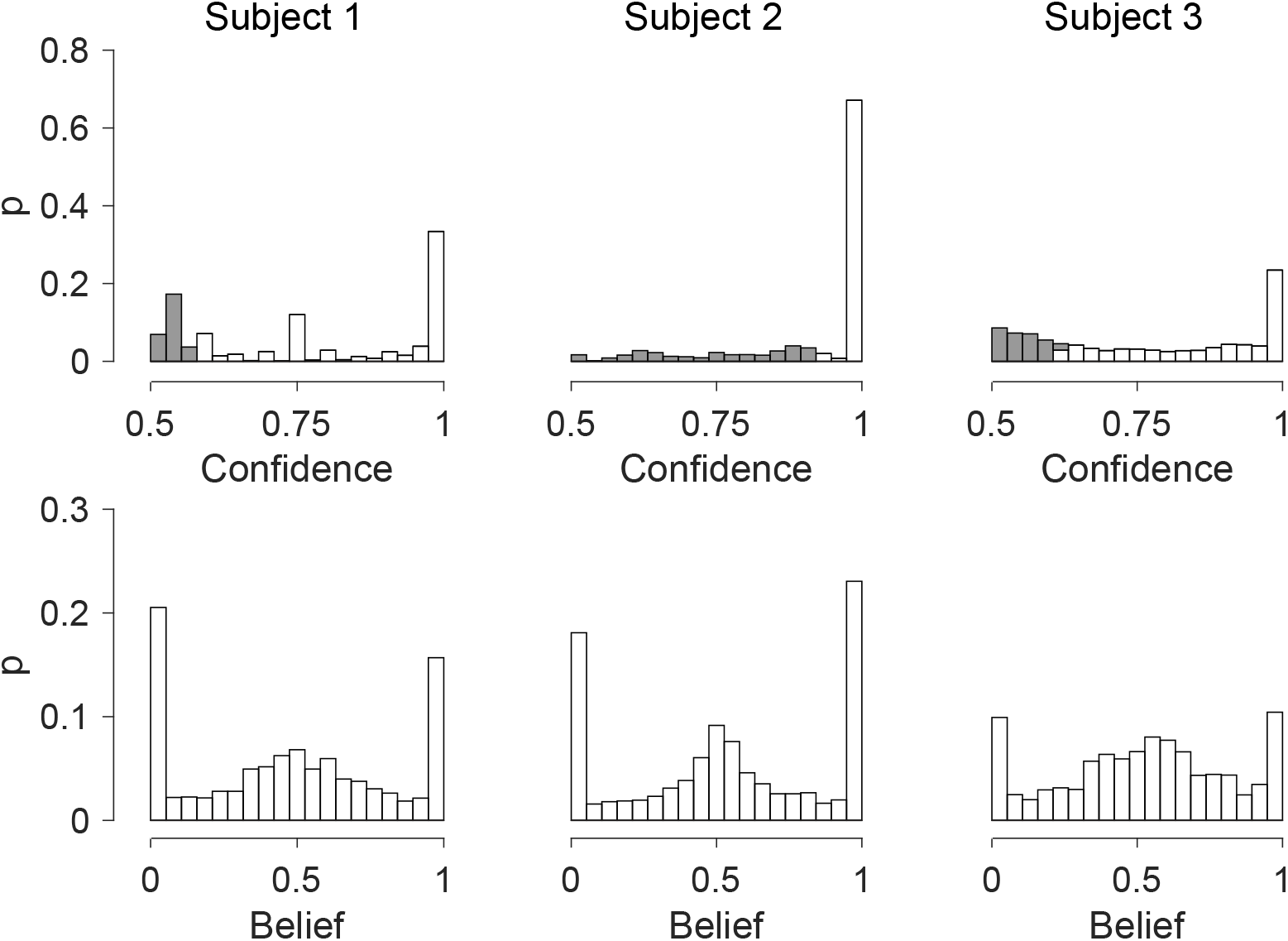
Distributions of confidence and belief reports. *Top row*, distributions of confidence reports for the 3 participants. Shading indicates the trials below the 30th percentile, which we designate “low confidence”. *Bottom row*, distributions of belief reports for the 3 participants.

**Figure S3.**
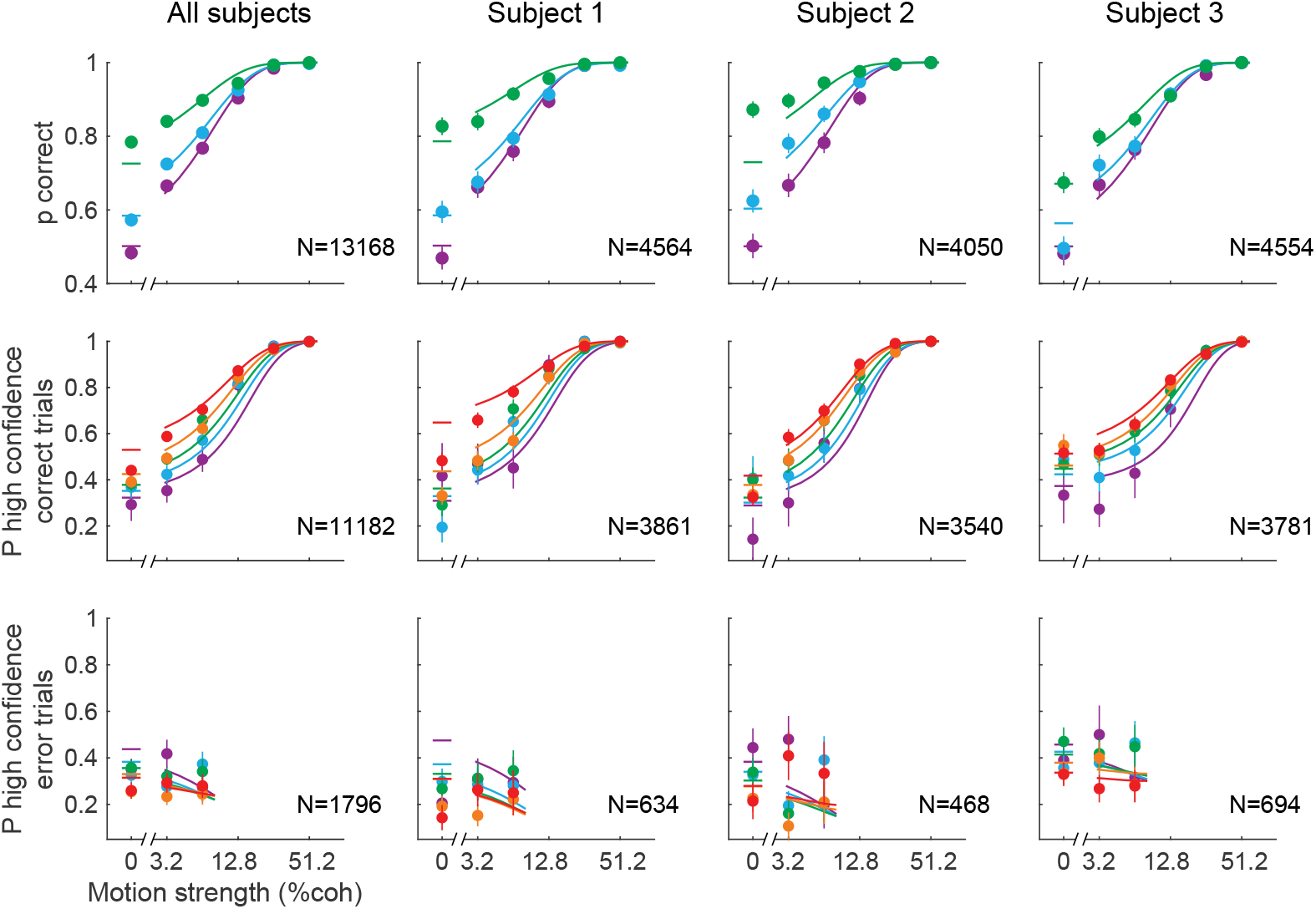
Fits of the Bayesian model to choice and confidence, combined and by subject. Left column shows combined data from all subjects, and columns 2-4 show the fits for the individual subjects. *Top*, Combined data is a reproduction of Figure 4A. *Middle*, Combined data is a reproduction of Figure 4D. *Bottom*, Same middle row, but for the error trials. Fits (solid lines) are fits using all trials, so dominated by the correct choices (errors constitute 13-17% of trials). Points comprising less than 10 trials are not shown.

**Figure S4.**
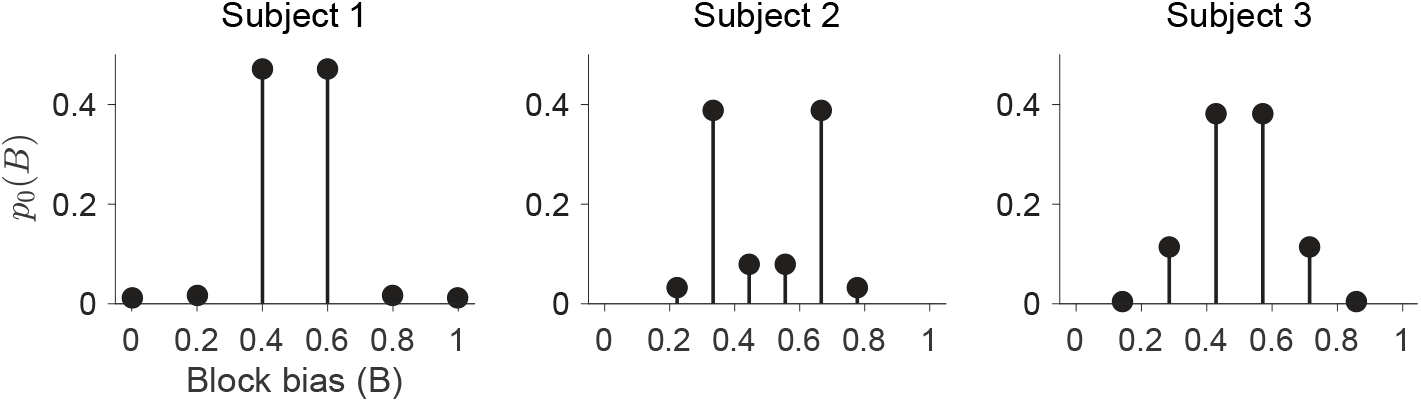
Initial prior probability distributions of base rate, *p*_0_(*B*). The graphs shows the probability distributions over base rate at the beginning of each block. They are obtained from the model fit to the choice and confidence. The base rate for the block is chosen from a uniform distribution over the six values *B* ∈ [0, 0.2, 0.4, 0.6, 0.8, 1]. However, the model allows for the possibility that the participant does not represent this prior veridically. In all cases, participants assigned higher probability to bias values close to ½.

